# Correct regionalisation of a tissue primordium is essential for coordinated morphogenesis

**DOI:** 10.1101/2020.08.29.273219

**Authors:** Yara E. Sanchez-Corrales, Guy Blanchard, Katja Röper

## Abstract

During organ development, tubular organs often form from flat epithelial primordia. In the placodes of the forming tubes of the salivary glands in the *Drosophila* embryo, we previously identified spatially defined cell behaviours of cell wedging, tilting and cell intercalation that are key to the initial stages of tube formation. Here we address what the requirements are that ensure the continuous formation of a narrow symmetrical tube from an initially asymmetrical primordium whilst overall tissue geometry is constantly changing. We are using live-imaging and quantitative methods to compare wild-type and mutants that either show disrupted cell behaviours or an initial symmetrical placode organisation, with both resulting in severe impairment of the invagination. We find that early transcriptional patterning of key morphogenetic transcription factors drives the selective activation of downstream morphogenetic modules, such as GPCR signalling that activates apical-medial actomyosin activity to drive cell wedging at the future asymmetrically-placed invagination point. Over time, transcription of key factors expands across the rest of the placode and cells switch their behaviour from predominantly intercalating to predominantly apically constricting as their position approaches the invagination pit. Misplacement or enlargement of the initial invagination pit leads to early problems in cell behaviours that eventually result in a defective organ shape. Our work illustrates that the dynamic patterning of the expression of transcription factors and downstream morphogenetic effectors ensures positionally fixed areas of cell behaviour with regards to the invagination point. This patterning in combination with the asymmetric geometrical set-up ensures functional organ formation.

## Introduction

Complex three-dimensional organs arise from simple tissue primordia, and in many cases these primordia are flat polarised epithelial sheets. Early in development the expression of the first patterning genes broadly sets up embryonic regions. This is followed by the activation of gene regulatory networks that specify the fate of tissue primordia in defined locations (1, 2). The patterning and fate determination gene products do not directly affect morphogenetic changes, but rather instruct the expression of downstream morphogenetic effectors that drive a tissue primordium down a path of defined physical changes. In many well-studied cases of tissue morphogenesis, such as mesoderm invagination or germband extension in the fly, many morphogenetic effectors are induced evenly across the tissue primordium. Regional differences in the physical changes or ‘behaviours’ of cells observed in these primordia can arise due to physical effects and feed-back or interference either from cells within the primordium or from surrounding tissues (3–5). A further layer of control and complexity is added as the primordia themselves can also be patterned to guide the differential behaviour and changes of groups of cells within a primordium. Such prepatterning of morphogenetic events within a single tissue primordium is much less understood.

We use the formation of a narrow-lumen tube in the *Drosophila* embryo as a model system to identify key requirements for successful organ formation. The symmetrical tubes of the embryonic salivary glands form from a flat, nearly circular epithelial primordium, the salivary gland placode (Fig. 1A). We recently uncovered a patterning of key cell behaviours that drive the initiation and earliest stages of the tube budding process (6). Interestingly, the point of invagination from the circular placode primordium is not in the centre of the placode, but in an asymmetric, eccentric position in the dorsal-posterior corner (Fig. 1A). Using quantitative morphometric methods we showed that the early apical constriction and associated cell wedging at the position of the invagination pit initiates the tissue bending (6, 7). Isotropic constriction driven by apical-medial actomyosin is concentrated near the forming invagination pit, but at a distance to the pit a second cell behaviour dominates: cell intercalation. Driven by a polarised junctional accumulation of actomyosin, cell intercalation events such as T1 exchanges and rosette formation and resolution help to elongate the tissue radially towards the invagination pit and contract it circumferentially, via circumferential neighbour gains (Fig. 1A’) (6).

**Figure 1.**
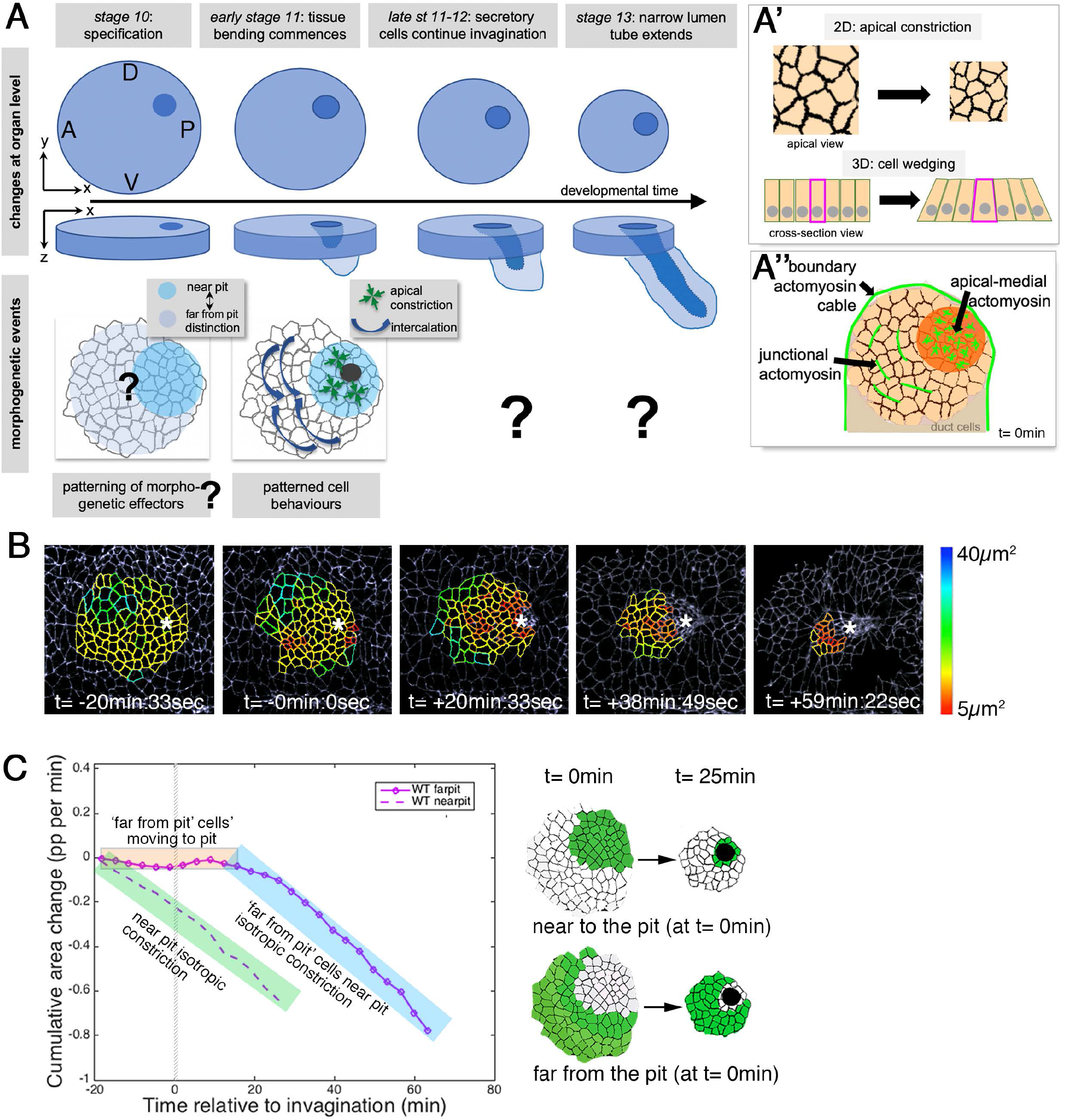
Patterned apical constriction remains fixed around the pit over time. **A** The tubes of the salivary glands form from a flat epithelial placode with a circular geometry. Cells invaginate through an asymmetrically positioned invagination pit at the dorsal-posterior corner, embryonic axes of anterior-posterior (AP) and dorso-ventral (DV) are indicated. Cell behaviours such as apical constriction and cell intercalation are highly patterned in the placode at early stages of tissue bending, and differ between cells near the pit and away from the pit (6). **A’** Apical constriction within the apical domain of an epithelial cell, a 2D change, equates to a behaviour of cell wedging within the 3D context of a whole cell, as shown in a cross-sectional view. A” Patterned cell behaviours are driven by distinct pools of apical actomyosin. Three pools of actomyosin can be distinguished in the placode: apical-medial actomyosin in cells near the invagination pit (dark orange), leading to isotropic apical constriction; polarised junctional actomyosin, driving the initiation of directed cell intercalation events in cells further away from the pit (light orange); a circumferential actomyosin cable at the boundary of the placode. **B** Stills of a representative segmented time-lapse movie, colour-coded for apical cell area size, showing the continued apical constriction of cells located near the invagination pit. See Supplemental Movie 1. **C** Analysis of apical area change pooled from 7 time-lapse movies for cells that were specified at t= 0min to be located near the invagination pit (dashed line; see schematic on the right) and cells that were specified at t= 0min to be located far from the invagination pit (solid line; see schematic on the right). Note that cells near the pit continuously constrict until internalised (green shaded box), whereas cells far from pit do not constrict significantly until t= 20min whilst moving closer to the pit (orange shaded box), but once they are in close proximity to the pit they also constrict until internalised (blue shaded box). Details of cell numbers and lengths of movies analysed are shown in Supplemental Figure 1.

This asymmetric set-up of the placode raises two questions: firstly, is the asymmetry required for successful organ formation or could a symmetrical set-up not lead to the same result? An indication that the asymmetry of the salivary gland placode is important for wildtype tube formation is given by the reports that in certain mutant situations where a more central invagination appears to form, the overall morphogenesis is disrupted and the invaginated structure that forms is sack-like and irregular. This is true for embryos lacking the transcription factor Hkb that is expressed early in the placode (8, 9), and could therefore potentially implicate Hkb in establishing early placode patterning.

Secondly, how is this asymmetry of the placode established prior to morphogenesis commencing and maintained throughout the process? Previous studies have shown that several proteins known to be expressed in the placode can be detected at the mRNA level initially in the dorsal posterior region of the placode. These include the transcription factor Forkhead (Fkh) (10), the dynein-associated protein Klar (9), as well as the kinase Btk29 (11). Defined expression of these and other factors at the dorsal-posterior corner could set up the initiation of cell shape changes in this region. In agreement with this, we previously found that *fkh^-/-^* mutants specifically fail to apically constrict in the dorsal-posterior corner, though they do show intercalation behaviour, and these mutants do not invaginate a tube (6). Following on from the early regionalised behaviours that drive the initial tissue bending and tube invagination (6), it is thus far unclear what mechanisms continue to drive the sustained invagination until all secretory cells have internalised. This could be due to continued transcriptional patterning and activation of downstream effectors, and/or could be due to mechanical signalling and feedback.

Here, we investigate how the asymmetry is set-up and maintained during salivary gland tube formation, in particular beyond the initial stages. We found that apical constriction is always strongest near the invagination pit at any moment in time. As more and more coronae of cells approach the region near the pit, they switch behaviour from predominantly intercalating to predominantly apically constricting/wedging. This continued switch to apical constriction leads to a smooth continued invagination process akin to a ‘standing wave’ of apical constriction, through which the placodal cells flow. The continued near-pit constriction is driven by apical-medial myosin, whose peak intensity and peak pulsatile strength is tracking in a fixed position close to the pit. Using live-imaging of fluorescently-tagged versions of Hkb and Fkh, we show that their expression levels pre-patterns where initial constriction is occurring. Both are upstream of a dynamic patterning of the GPCR ligand Fog, that in turn activates apical-medial myosin. Loss of the correct pre-patterning, such as in *hkb^-/-^* mutants or when Fog is overexpressed, leads to loss of the symmetrical final organ shape with a narrow lumen and instead gives rise to expanded and sack-like glands with widened lumena.

Our work establishes that after the tissue is patterned at the posterior corner, cells follow dynamic rules of behaviour as they approach the pit. Symmetrical tube morphogenesis thus relies on a regionalisation pattern of the primordium long before cells show distinctive behaviours. Establishing general mechanisms that modulate the orderly and repetitive behaviour of cells to form correctly shaped simple model organs such as the salivary glands will assist the understanding morphogenesis of other more complex tubular organs in mammals.

## Results

### Patterned apical constriction remains fixed around the pit over time

The initial morphogenetic change that occurs during the budding of the tube of the salivary glands in the *Drosophila* embryos is the apical constriction of cells at the point of the future site of invagination, the pit (Fig. 1A) (6, 8, 10, 12). We recently provided the first quantitative morphometric analysis of the early stages of this process, and revealed that firstly isotropic apical constriction near the future pit commences long before changes at the tissue level are apparent, and that secondly the apical constriction is indicative of a 3D cell behaviour of wedging of cells at the pit (Fig. 1A’). Importantly, we uncovered a strong regionalisation of cell behaviours during the early stages of tube budding: cells near the invagination point predominantly showed wedging behaviour while cells at a distance from the pit predominantly showed intercalation behaviour (6)(Fig. 1A). What is unclear is how the early tube initiation via isotropic constriction and wedging, combined with directional intercalation in cells away from the forming pit, evolves into the continuous invagination and formation of a symmetrical narrow-lumen tube from the asymmetrically placed invagination pit. In one scenario, cell behaviour is fixed at early stages of budding, i.e. cell wedging is restricted to the pit and only apparent during early stages. In a second scenario, cell behaviour depends on a cell’s position relative to the pit and thus cell behaviours change as cells move towards the invagination point.

To distinguish between these scenarios, we segmented and tracked placodal cells from time-lapse movies covering a −40 min to +70 min time interval relative to the first occurrence of tissue bending during the invagination process (see Suppl. Fig. S1 for time intervals of analysed movies and cell numbers analysed) and performed quantitative analyses (Fig. 1B, C). At the latest time point covered, the majority of secretory cells had completely internalised and formed the internal tube. Over the whole period analysed, cells positioned near the invagination pit always displayed constricted apices (Fig. 1B, C and Supplemental Movie 1), suggesting that cells changed their behaviour depending where they were positioned in the placode with respect to the invagination point. Cells near the pit displayed isotropic constriction and thus a negative cumulative apical area change from the start (Fig. 1C, green highlighted), whereas cell far from the pit did not change their apical area over the initial ~20 min (Fig. 1C, orange highlighted). Cells were defined as ‘far from the pit’ at time point t= 0min. These ‘far from the pit’ cells then moved into a position near the pit through cell intercalations and due to previous cells at the pit internalising. Once these originally ‘far from the pit’ cells were in such ‘near the pit’ position they also displayed isotropic apical constriction and thus negative apical area change (Fig. 1C, blue highlighted).

These data strongly suggest that an initially prepatterned behaviour, isotropic constriction at the pit and intercalation further away, is dynamically adjusted during tube budding from a flat epithelial sheet, so that cells within a similar distance to the invagination pit display the same behaviour

### Sustained apical constriction near the pit is driven by sustained apical-medial myosin

Early apical constriction at the pit is driven by a highly dynamic apical-medial actomyosin network (6, 7). We next sought to determine whether the apical-medial actomyosin dynamics would differ among regions in the placode. We analysed apical-medial actomyosin levels in cells across the placode, and firstly illustrate our findings in two snapshots: cells near the future invagination point at t = 0min showed strong accumulation of apical-medial myosin (Fig. 2A, A’’, C; see also Supplemental Movie 2), whereas cells far from the pit and close to the anterior placode boundary (Fig. 2A, A’) showed only junctional myosin II accumulation. At t = 30 min the cells now located right next to the invagination pit continued to displayed strong apical-medial myosin accumulation (Fig. 2B, B’’), but in addition former ‘far from the pit’ cells near the anterior boundary that were now in a closer proximity to the pit also started to display apical-medial actomyosin (Fig. 2B, B’). Thus, being moved into a position near the invagination pit appeared to determine levels of apical medial myosin accumulation. Looking placode-wide over the whole time period analysed, the peak of the apical-medial myosin fluorescence intensity tracked in a near-fixed position to the edge of the pit over time (Fig. 2D). The location of the highest medial myosin intensity was also where cells showed the shortest cycle lengths of medial myosin II fluctuations (Fig. 2E). Interestingly, cells near the pit at t = 0 min showed overall shorter myosin II cycle length (the time period elapsed between two peaks of highest myosin II intensity during the oscillation, see schematic in Fig. 2E) compared to cells far from the pit (Fig. 2F). As longer cycle length correlates with unproductive myosin cycles (7), this was in line with cells away from the pit showing less apical constriction. Furthermore, the strength of myosin oscillations can be expressed as the product of the amplitude and frequency of the oscillation (7). This myosin strength across the placode at t = 0 min was much higher in cells near the invagination point when plotted against the distance to the pit, with a gradual decrease in myosin strength towards the placode boundary (Fig. 2G). Cell defined as ‘far from the pit’ at t = 0 min though increased their myosin strength over time (and concomitantly decreased their cycle length; see Supplemental Figure S2), so that by t = 55 min their myosin strength had increased to levels previously shown by ‘near the pit’ cells at t = 0 min (Fig. 2H).

**Figure 2.**
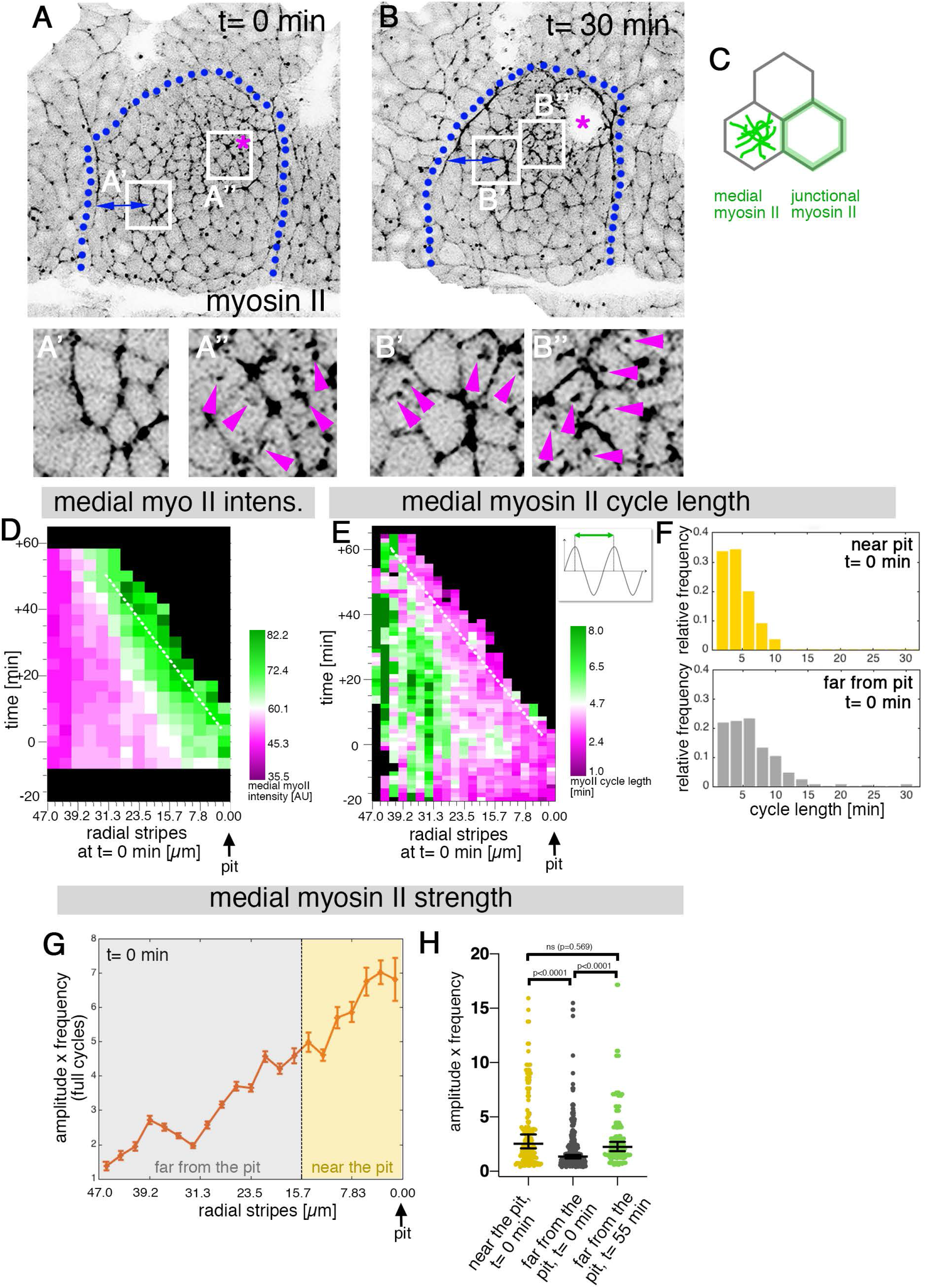
Sustained apical constriction near the pit is driven by apical-medial myosin. **A-B”** Still images of a time-lapse movie of embryos expressing SqhGFP to label myosin II at t= 0min (**A-A”**) and t= 30min (**B-B”**). White boxes indicate the corresponding higher magnifications shown below, with **A”,B”** showing cells that are at these timepoints located near the pit, and **A’,B’** showing cells near the boundary (small blue double arrows indicate similar closeness to the boundary). Magenta asterisk marks the invagination pit, blue dotted lines mark the boundary of the placode. **C** Schematic apical view of epithelial cells, illustrating junctional and apical-medial pools of actomyosin. Arrows in **A’’**, **B’**, **B’’** point to apical-medial actomyosin. See also Supplemental Movie 2. **D** Spatial representation of the average medial myosin intensity from an exemplary time lapse movie, with radial location (collapsed into stripes) of cells specified at t= 0min. The dashed white line marks the peak of medial myosin intensity that is always adjacent to the invagination pit as cells flow into it over time. **E** Spatial representation of the distribution of the myosin cycle length across the salivary gland placode, with radial location (collapsed into stripes) of cells specified at t= 0min. The dashed line marks the region of shortest cycle length that is always adjacent to the invagination pit as cells flow into it over time. The mean of 3 movies is shown. The schematic illustrates one cycle of a myosin pulsation defined as periodic increases and decreases in medial myosin II intensity. **F** The distribution of myosin II cycle lengths between cells near the pit and far from the pit varies: the median of the cycle length for cells near the pit is 3.7 min (n =146) and SD+/- 2.08 min, while cells far from the pit have a median myosin II cycle length of 4.64 min (n=301) and SD+/- 2.72 min. Thus, the cells that are located far from the pit show longer cycle length (p < 0.0001 Mann Whitney test). Data is pooled from 3 movies at t=0. **G** Spatial representation of the strength of myosin oscillations in radial stripes from the pit (0 μm) to the boundary of the placode (47μm) at t= 0min. Mean and standard error of the mean are shown. Data are pooled from 3 movies ranging from −7.5-56.25 min; −16.76 to 55.48 min and −17.42 to 67.26min. Regions corresponding to cells near the pit and far from the pit are indicated by coloured shading. **H** Medial myosin strength, expressed as the product of amplitude x frequency of the oscillations, for all cells analysed from 3 movies, split into cells located near to the pit (n=148) compared to cell located far from the pit (n=302) at t=0 from 3 movies. Cells near the pit have significantly higher medial myosin strength than cells located far from the pit; statistical significance as determined (Mann Whitney test, with p<0.0001). Cells far from the pit at later time points (t=55-60 min, n = 134) increased the medial myosin strength and were not significantly different from cells near the pit at t=0 (Mann Whitney test, p = 0.5690). See also Supplemental Figure 2 and Supplemental Movie 2.

Thus, our data show that the early asymmetric set-up of high apical-medial myosin II intensity and activity clustered near the pit was maintained during the continued invagination of the tube, in turn driving the continued apical constriction near the pit.

### *hkb^-/-^* mutants show a delayed symmetrical apical constriction

In order to uncover the role of the initial asymmetric patterning of cell behaviours within the placode, we turned our attention to mutant situations: we already knew that Fkh was important, as *fkh^-/-^* mutants do not show localised apical constriction or any invagination (6). A possibly more interesting situation is present in a previously published mutant in the transcription factor Huckebein (Hkb) that was reported to show a central invagination pit, combined with malformed invaginated salivary glands at later stages (8).

We collected and segmented time-lapse movies of *hkb^-/-^* mutant embryos over the same time period as the control wild-type embryos (Fig. 3). Whereas in the wild-type, at t~ +40min, most secretory cells of the salivary gland placode have already invaginated (Fig. 3 A’’’, B’’’, E’’’), in the *hkb^-/-^* mutants no invagination had occurred yet (Fig. 3 C’’’, D’’’, F’’’). Eventually, these embryos invaginated a dilated tube through an enlarged pit (see Supplemental Figure S3_2 and Supplemental Movies 5 and 6). Analysing apical-medial actomyosin intensity and distribution in the wild-type compared to *hkb^-/-^* embryos, we found that the striking clustered accumulation of apical-medial actomyosin near the future invagination pit at t= 0min in the wild-type (Fig. 3B’) was completely absent in *hkb^-/-^* embryos (Fig. 3D’’). Instead, over the time period analysed here only at t~ +40min a group of cells in the centre of the placode of *hkb^-/-^* mutant embryos displayed constricted apices (Fig. 3D’’’), inducing a shallow tissue bending in this region (see also Suppl. Fig. S3_2).

**Figure 3.**
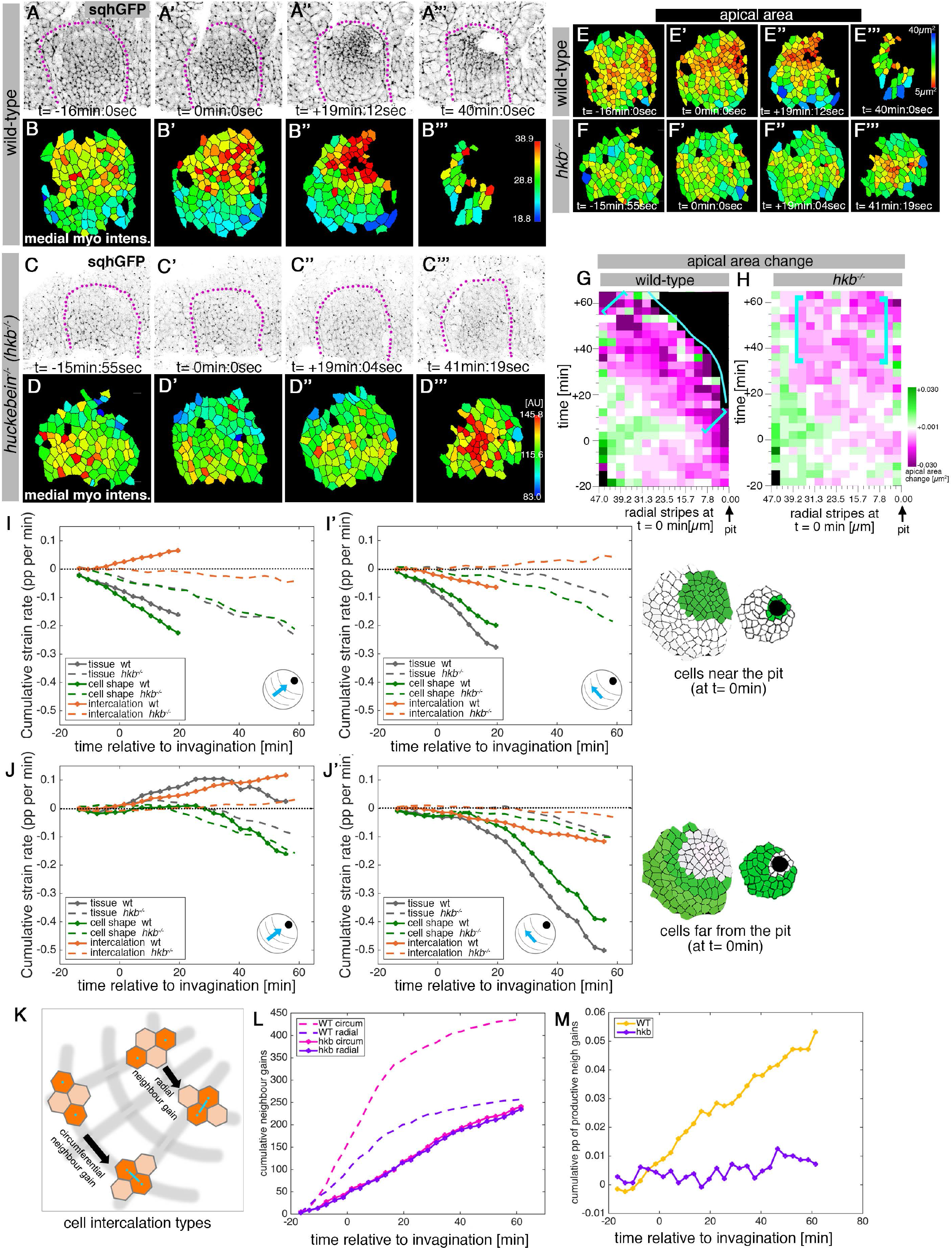
hkb^-/-^ mutants show a delayed symmetrical apical constriction. **A-D”’** Stills of a representative wild-type (**A-B”’**) and a *hkb^-/-^* mutant (**C-D”’**) embryo time lapse movie at the indicated time points. **A-A”’** and **C-C”’** show the SqhGFP channel of the movie visualising myosin II, **B-B”’** and **D-D”’** show the average apical-medial myosin II intensity of segmented placodal cells. *hkb^-/-^* mutants show apical-medial accumulation of myosin II at the centre at a delayed time point. Indicated colour scales correspond to the 5-95% range in each movie. Magenta dotted lines indicate the boundary of the placode. **E-F’’’** The delayed accumulation of myosin II in *hkb^-/-^* mutants is mirrored by the changed the apical constriction pattern, with the wild-type constricting in the dorsal-posterior corner (**E-E”’**) and *hkb^-/-^* mutants in the centre, but very delayed (**F-F”’**). Colour scales for cell areas as in Figure 1. **G,H** Rate of apical area change of cells in wild-type (**G**; data from 7 movies) and *hkb^-/-^* mutant (**H**; data from 7 movies) embryos shown over time and across radial distance of cells from the pit location (0 μm) at t = 0 min. 47 μm represents the boundary of the placode. The blue line in **G** indicates where cells have just disappeared into the pit in the wild-type. Greatest apical area change in the wild-type is always confined to the area in front of the pit (brackets in **G**), whereas in *hkb^-/-^* mutant embryos cells in a broader central region show a delayed apical constriction (brackets in **H**). See also Supplemental Figure 3_1. **I-J’** Regional breakdown of time-resolved cumulative strain rates, with regions defined at t= 0min based on wild-type geometry. For cells ‘near the pit’ in the wild-type, tissue constriction dominates (solid grey curves in **I**, **I’**) and is due to isotropic cell constriction (solid green curves in **I, I’**), whilst intercalation only plays a minor role in this region (solid orange curves in **I, I’**). Cells in this region have completely internalised by about t= 20min. By contrast, in *hkb^-/-^* mutant embryos, cells ‘near the pit’, i.e. near the predicted pit position, show strongly reduced tissue (dashed grey curves in **I, I’**) and cell strain rates (dashed green curves in **I, I’**) and nearly no intercalation (dashed orange curves in **I, I’**). For cells ‘far from the pit’ in the wild-type, the tissue elongates towards the pit until t= 20min (solid grey curve in **J**), with a corresponding contraction circumferentially (solid grey curve in **J’**), and this is predominantly due to cell intercalation (solid orange curves in **J, J’**). Beyond t= 20min, these cells have reached the invagination pit and also constrict isotropically, thereby leading the tissue change (solid grey curves in **J, J’** >20min) to mirror the cell shape change (solid green curves in **J, J’** >20min). By contrast, in *hkb^-/-^* mutant embryos, cells ‘far from the [predicted] pit’ show no change of tissue, cell shape or intercalation strain rates prior to 20min (dashed curves in **J, J’**), and after this point show slight isotropic constriction, reflecting the observed centrally located constricted apices (**F’’’**). The corresponding instantaneous strain rate plots can be found in Supplemental Figure 3_3. Data from 9 wild type movies and 5 *hkb^-/-^* movies were analysed (see Supplemental Figure 3_1). **K-M** Quantification of neighbour gains as a measure of T1 and intercalation events. Examples of a circumferential neighbour gain (leading to radial tissue expansion), and a radial neighbour gain (leading to circumferential tissue expansion) are shown in **K**. **L** Circumferential neighbour gains dominate over radial neighbour gains in the wild-type (dashed curves), with the rate of neighbour exchanges dropping beyond 20min. In contrast, in *hkb^-/-^* mutant embryos the amount of circumferential and radial gains is identical (solid curves). **M** Rate of productive gains, defined as the amount of circumferential neighbour gains leading to radial tissue elongation and expressed as a proportion (pp) of cell-cell interfaces tracked at each time point. The cumulative proportion of productive neighbour gains is significantly different between wild-type and *hkb^-/-^* mutants (two-sample Kolmogorov- Smirnov-test, p<0.05). Data pooled from 9 wild type and 5 *hkb^-/-^* mutant movies. See also Supplemental Figure 3_1.

The apical area analysis strongly suggested that the initial asymmetrical set-up of cell behaviours in the placode might be disrupted. We therefore performed a strain (deformation) rate analysis of *hkb^-/-^* mutants in comparison to wild-type embryos to assess this quantitatively. The strain rate analysis is based on the assumption that any change in tissue shape can be accounted for by changes in cell shape and cell intercalations (6, 13). In the wild-type, cells that were located near the invagination pit (defined at t = 0 min) constricted isotropically, with near equal change in strain at the tissue and cell shape level, and very little intercalation strain (Fig. 3I, I’, solid curves). These cells in the wild-type could only be tracked up to t = 20 min as at this point they had all invaginated into the embryo to form the initial part of the tube. In contrast, in the *hkb^-/-^* mutants cells specified at t = 0 min to be near the coordinates of the position of where the invagination pit would be in the wild-type showed a much reduced tissue and cell shape strain rate, with near identical values in both radial and circumferential orientation (Fig. 3I,I’, dashed curves; see also Suppl. Fig. S3_3). For cells that at t = 0 min were located away from the pit, in the wild-type we observed signatures of convergence and extension through cell intercalations within the first approximately 20 min after invagination commenced, as also reported previously (Fig. 3 J, J’, solid curves; tissue and intercalation strain rate showing expansion radially and contraction circumferentially; (6)). Interestingly, we then observed a clear switch in cell behaviour to a second phase after t = +20 min. Once these cells were located closer to the pit position, they now displayed near isotropic contraction (beyond t = +20 min). In the *hkb^-/-^* mutants, for cells marked as far from the theoretical pit position at t = 0 min, there was nearly no strain rate observed at either tissue, cell shape or intercalation level initially until t ~ +20 min, after which the cells showed a low level of isotropic constriction (Fig. 3J, J’, dashed curves).This constriction corresponded to the centrally positioned constricting apices described above in Fig. 3D’’’ and F’’’. There was essentially zero intercalation strain rate observed during the period recorded in the *hkb^-/-^* mutants (Fig. 3J, J’, orange dashed curves).

To assess whether the low or absent strain rates indicated a very static tissue we analysed individual events of neighbour gains across the tissue in wild-type and *hkb^-/-^* mutant embryos. We compared two types of intercalation events, those leading to circumferential neighbour gains that in the wild-type explain the observed intercalation strain rates, and those leading to radial neighbour gains, and thus potentially opposing the changes seen at the tissue and intercalation strain rate level (Fig. 3 K). In the wild-type, circumferential neighbour gains outweigh radial ones by a large margin throughout (Fig. 3L), leading to a steady increase in cumulative productive neighbour gains (Fig. 3M). In contrast, in *hkb^-/-^* mutant embryos circumferential and radial neighbour gains are overall much reduced in number and occur with equal frequency (Fig. 3L), thus leading to near zero productive neighbour gains (Fig. 3M).

These analyses show that in the absence of Hkb, the initial radial patterning of cell behaviours across the placode is lost, and cells located at a position ‘near the pit’ in the *hkb^-/-^* mutants clearly fail to apically constrict, suggesting that Hkb could be involved in the specification of the pit-behaviour observed in wild type.

### The salivary gland placodal primordium is asymmetrically patterned by Hkb and Fkh prior to morphogenesis

The loss of correct patterning of cell behaviours in the early placode in *hkb^-/-^* mutant embryos suggested that Hkb was involved in establishing these in the wild-type. Furthermore, *hkb* expression has been reported to show a dynamic pattern by *in situ* hybridisation and β-gal reporter labeling (8). In order to be able to analyse Hkb protein levels dynamically and in comparison to cell shapes, we generated a Venus-tagged version of Hkb using CRISPR/Cas9 to tag the endogenous protein. Using this *in vivo* reporter, we detected Venus-Hkb at high levels concentrated in the dorsal-posterior region of the just forming placode, in the area where the invagination pit will form, already at t = −30min prior to the first tissue bending (Fig. 4 A, A’, D, M and Suppl. Fig. S4B-B’’), and even as early as t = −63 min prior to invagination (Suppl. Fig. S4A). The expression then expanded more broadly and anteriorly across the whole placode over time (Fig. 4B-C’, E, F, M and Supplemental Movie 7). At the moment of first tissue bending (t = 0 min) Venus-Hkb was found across the placode, with elevated levels still at the invagination pit position, and a second increase towards the anterior. At late stages (t >> 0 min) Venus-Hkb levels close to the invagination point were lower than further anterior (Fig. 4M).

**Figure 4.**
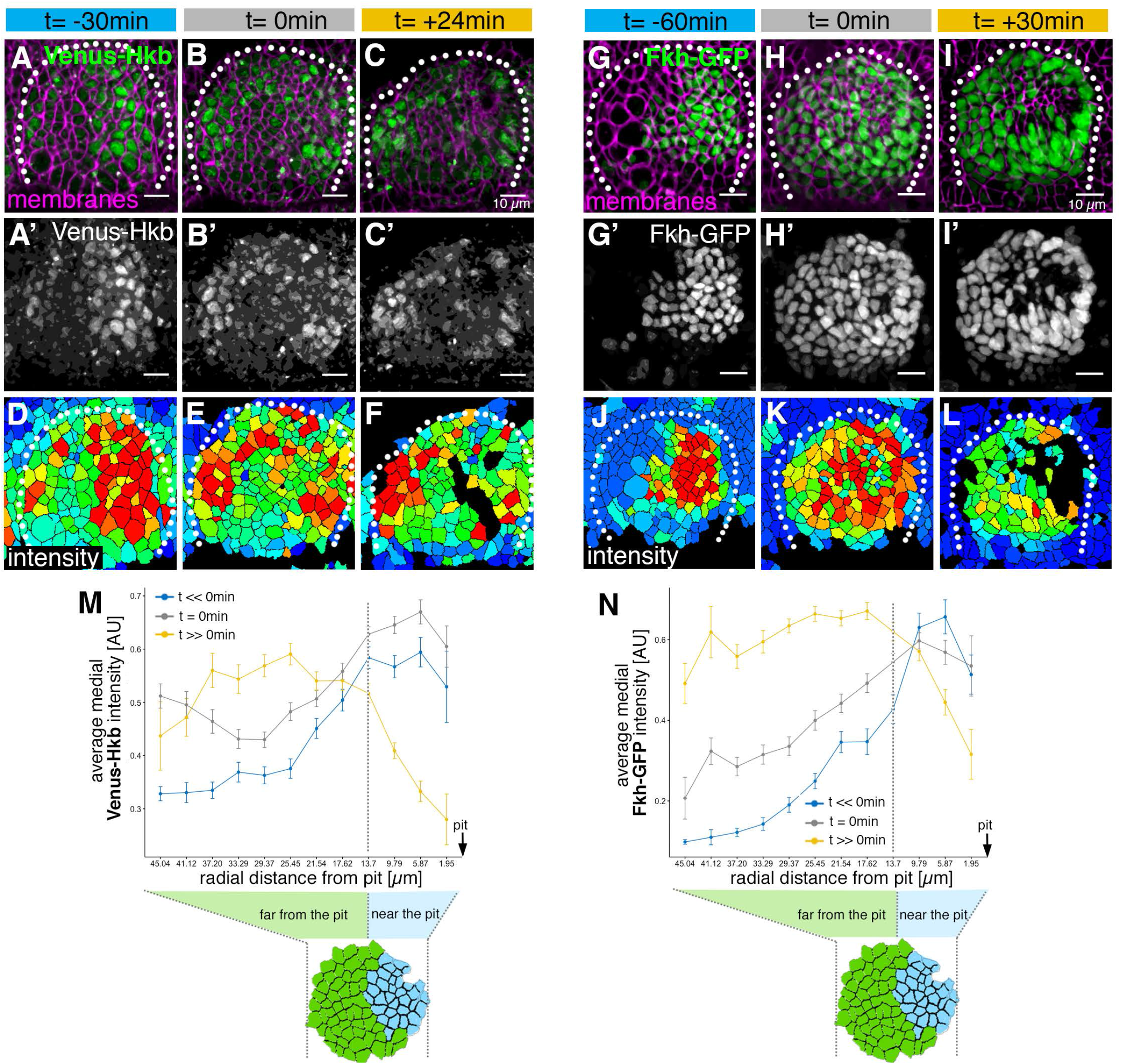
The salivary gland placodal primordium is asymmetrically patterned by Hkb and Fkh prior to morphogenesis. **A-F** Stills of a time lapse movie of embryos with endogenously tagged Hkb, Venus-Hkb, at t= −38min, +0min and +38min show the dynamic pattern of expression of Hkb. Expression starts at the posterior corner before tissue bending commences. Cell membranes are in magenta in **A-C**, Venus-Hkb is green in **A-C** and as a single channel in **A’-C’**. **D-F** show the quantification of Venus-Hkb fluorescence intensity at these time points. **G-L** Stills of a time lapse movie of embryos with Fkh tagged by GFP under endogenous expression control, at t= −60min, +0min and +30min show that as early as one hour before tissue bending, Fkh is already expressed at the posterior corner. Cell membranes are in magenta in **G-I**, Fkh-GFP is green in **G-I** and as a single channel in **G’-I’**. **J-L** show the quantification of Fkh-GFP fluorescence intensity at these time points. **M** Venus-Hkb expression starts at the posterior corner in a region similar to ‘near the pit’ population (between pit location and dashed line). At the start of invagination (around t= 0min) the expression increases across the placode, in particular at the anterior edge. At late stages, the level of Venus-Hkb expression decreases in the pit region, but it remains high in the rest of the placode. Data is pooled from selected frames of time-lapse movies and corresponding fixed samples. Number of cells are as follows: t<<0 min (equivalent to stage 10), n = 777 cells from 8 embryos; t=0 min (equivalent to early stage 11), n = 1186 cells from 10 embryos and t>>0 min (equivalent to late stage 11/early stage 12), n = 885 cells from 10 embryos. **N** Fkh-GFP expression initiates at the posterior corner and increases to cover the whole placode over time. Data is pooled from selected frames of time-lapse movies and corresponding fixed samples. Number of cells are as follows: t<<0 min (equivalent to stage 10), n = 365 cells from 4 embryos; t=0 min (equivalent to early stage 11), n = 1026 cells from 9 embryos and t>>0 min (equivalent to late stage 11/early stage 12), n = 659 cells from 9 embryos.

As mentioned above, embryos mutant in Fkh also show a lack of early apical constriction at the dorsal-posterior corner of the placode (6, 10). Similar to Hkb, analysis of *fkh* mRNA and protein levels in fixed embryos indicated an early enrichment near the forming invagination pit (6, 10). Using an *in vivo* genomic GFP-Fkh reporter (14), we followed Fkh protein levels and patterning over the same time period whilst also assessing apical cell shapes (Fig. 4 G-L). Already at t= −60min prior to tissue bending, Fkh-GFP was visible in the cells near the future invagination pit (Fig. 4 G, G’, J, N and Suppl. Fig. S4 C-C’’), from where it spread anteriorly and ventrally across the whole placode (Supplemental Movie 8). At t = 0min, Fkh-GFP was already detected in all cells of the placode, though with levels still highest near the pit and decreasing towards the anterior edge of the placode (Fig. 4 H, H’, K, N). At t = +30 min and beyond, levels of Fkh-GFP were more uniform across the placode (Fig. 4 I, I’,L, N).

Thus, both Hkb and Fkh transcription factors showed a clear temporally and spatially graded pattern of expression within the early placode that developed dynamically. Changes were especially pronounced at the dorsal-posterior corner, with highest levels here already an hour before tissue bending commenced. Combined Hkb and Fkh action could thereby define the future position of the invagination pit. These dynamic changes could also trigger the graded and patterned expression of targets of both transcription factors, leading to the observed patterned cell behaviours we uncovered, as well as the sustained apical constriction tracking near the pit.

### Asymmetric Fog expression is controlled by Hkb and Fkh and is upstream of early differential behaviours in the placode

With the transcription factors Fkh and Hkb both displaying intriguing protein expression patterns across the early placode pre-morphogenesis, targets of both factors are likely to play key roles in instructing cell behaviours such as wedging and cell intercalation. Interestingly, an upstream activator of apical-medial actomyosin activity, the GPCR-ligand Folded gastrulation (Fog) was found to be to be dependent on Fkh (15, 16) (Fig. 5J) and suggested to be downstream of Hkb [(8); though as ‘data not shown’]. Fog is an apically secreted ligand that acts in an endocrine fashion, and thus the cells that switch on Fog expression will apically constrict (16). Previously published *in situ* hybridisation of *fog* mRNA suggested increased levels near the forming pit, similar to the early Hkb and Fkh enrichment (17). Fog expression and signalling in particular could not only be upstream of the isotropic constriction of cells near the forming pit, but also be involved in maintaining this behaviour in new coronas of cells moving near the pit over time.

**Figure 5.**
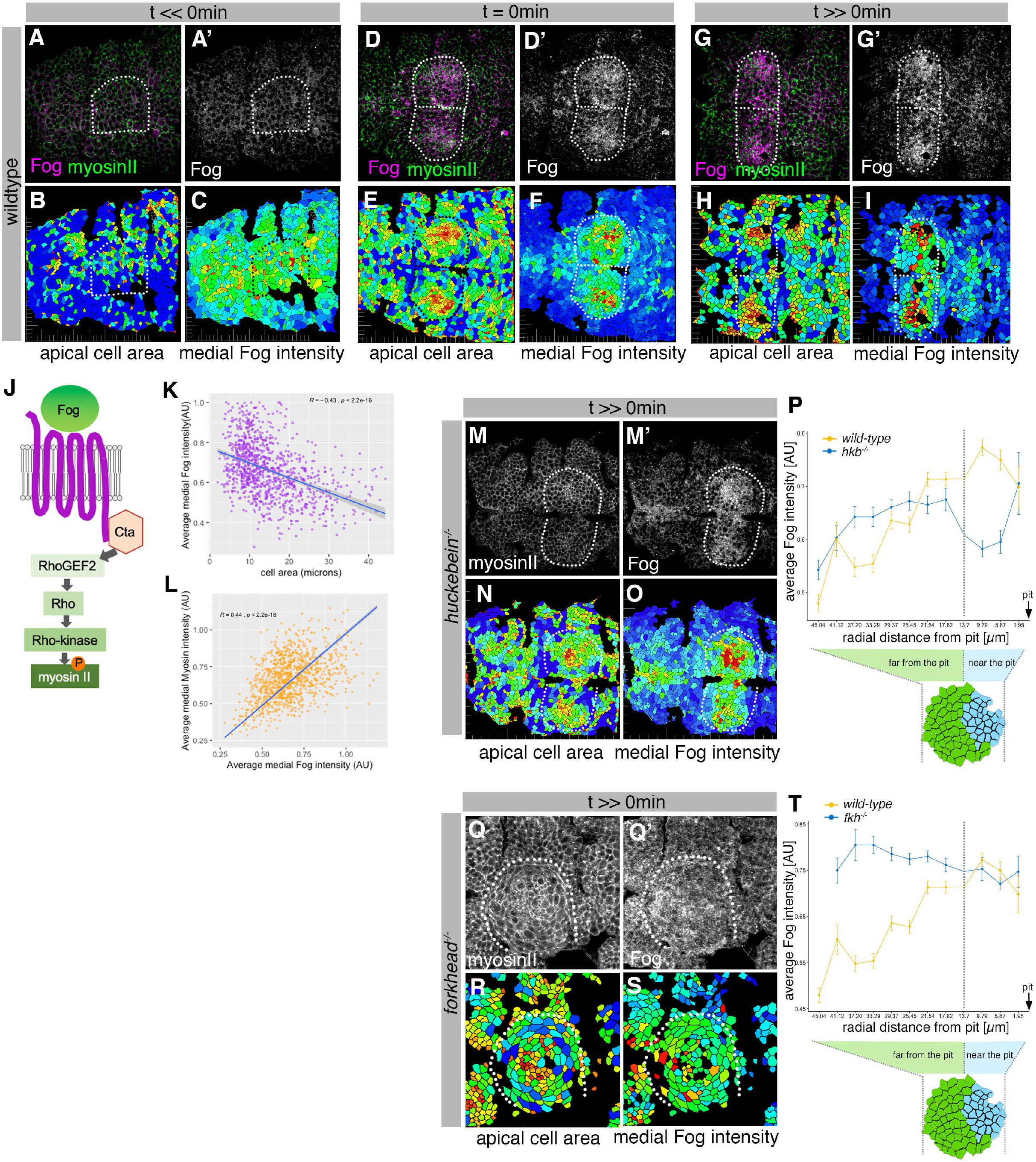
Asymmetric Fog expression is controlled by Hkb and Fkh upstream of early differential behaviours in the placode. **A-I** Wild-type embryos labelled for Fog protein (the ligand to the GPCR upstream of Rho- dependent myosin II activation, **J**) and myosin II, visualised by SqhGFP. Fog is magenta in **A, D, G** and as a single channel in **A’, D’, G’**, myosin II is green in **A, D, G**. Also shown are corresponding quantifications of apical cell area (**B, E, H**) and average medial Fog fluorescence intensity (**C, F, I**). Time points analysed were before any apical constriction and tissue bending commencing (t<< 0min; ~ stage 10), at t= 0 min (~early stage 11) and once invagination had commenced (t>>0 min; ~ late stage 11/early stage 12). **J** Schematic of the GPCR pathway leading to myosin II activation. **K** Average medial Fog fluorescence intensity negatively correlates with apical cell area. Pearson coefficient r = −0.436; n = 823 cells from 7 embryos. **L** Average medial myosin II fluorescence intensity positively correlates with average medial Fog fluorescence intensity. Pearson coefficient r = 0.439; n = 823 cells from 7 embryos. **M-O** In *hkb^-/-^* mutant embryos at a time point where invagination would have been well advanced in wildtype embryos (t>>0 min), medial myosin II (**M**) and Fog fluorescence (**M’** and quantified in **O**) are concentrated in cells in the centre of the placode, where cells also show constricted apices (**N**). **P** Quantification of average medial Fog fluorescence intensity according to radial position in wild-type versus *hkb^-/-^* mutant embryos. Whereas in the wild-type Fog intensity peaks at the posterior end of the placode where the invagination pit forms, in *hkb^-/-^* mutant embryos Fog is dramatically reduced at the invagination pit. Number of cells: wild-type n=823 from 7 embryos; *hkb^-/-^* n =765 cells from 5 embryos. **Q-S** In *fkh^-/-^* mutant embryos at a time point where invagination would have well advanced in wildtype embryos (t>>0min), medial myosin II (**Q**) and Fog fluorescence (**Q’** and quantified in **S**) are very homogenous across the placode and no longer enriched within the placode compared to the surrounding epidermis. Apical area quantification shows there is only a mild central cell constriction and no invagination (**R**). **T** Quantification of average medial Fog intensity in *fkh^-/-^* mutant embryos. In contrast to wildtype embryos with Fog enrichment at the posterior pit, levels of Fog are more homogenous in the placode in *fkh^-/-^* mutant embryos with an increase towards the anterior. Levels are comparable to the rest of the epidermis (see Supplemental Figure 5). The intensity curves shown for wild-type and *fkh^-/-^* mutant embryos are comparable in their shape, though the absolute intensities shown are not directly comparable, as fluorescence intensity was normalised in each image by dividing the average fluorescence per cell by the 98^th^ percentile value to account for embryo to embryo variability in staining efficiency. Number of cells: wild type n = 823 from 7 embryos; *fkh^-/-^* mutant n= 512 cells from 5 embryos.

We therefore decided to carefully analyse Fog protein levels in comparison to apical area and apical-medial myosin intensity in salivary gland placodes from early to late invagination stages (Fig. 5A-I). Fog levels were low at mid stage 10 concomitant with placode specification (Fig. 5A, A’, C). Just before tissue bending commenced at t= 0min, Fog was already strongly enriched in cells near the forming invagination pit, and these cells were the ones already showing decreased apical area (Fig. 5 D-F). Fog protein levels remained enriched near the invagination point once the tube had started to internalise (Fig. 5G-I). In fact, Fog protein levels in cell were positively correlated with both smaller apical area (Fig. 5K) as well as higher average apical-medial myosin II intensity (Fig. 5L; Suppl. Fig. S5).

*hkb^-/-^* mutant embryos, as detailed above, showed a very clear disruption of the early asymmetric placodal patterning. At late stage 11, Fog protein levels appeared to be similar to wild-type and enriched in the placode compared to the surrounding epidermis, though the pattern of Fog enrichment was altered. Cells at the posterior corner, where the invagination pit would have formed in the wild-type, lacked high Fog protein levels, whereas only cells in the centre of the placode now showed high levels of Fog protein (Fig. 5 M-O). When quantitatively analysing many *hkb^-/-^* mutant embryos in comparison to wild-type, the peak intensity of Fog protein was clearly shifted to a central, much more symmetrical, position (Fig. 5 P). In contrast, although we could detect some Fog protein in *fkh^-/-^* embryos at late stage 11, it was no longer enriched at the position where the invagination pit forms in the wild-type. The levels also did not appear raised anywhere within the placode compared to the surrounding epidermis (Fig. 5Q-S and Suppl. Fig. S5 D-D’’), as the quantification of many placodes confirmed (Fig. 5T). In both mutants, altered levels and patterning of Fog protein still correlated with the now altered levels and patterns of residual apical-medial myosin (Fig. M, Q).

Thus, the asymmetrical set up of the salivary gland placode prior to initiation of tube budding relies on a correct patterning of key transcripts factors, Hkb and Fkh. These factors are in turn upstream of GPCR/Fog signalling that leads to patterned and sustained apical-medial myosin activity and hence contractile cell behaviours.

### Changes to invagination pit size or position impact invagination and perturb final organ shape

Our data so far show that the asymmetrical invagination point depends on patterning by transcription factors and downstream GPCRs/Fog signalling at the posterior corner, leading to a focussed and point-like invagination pit. We next sought to test the importance of this asymmetric set-up for successful morphogenesis of a narrow-lumen and symmetrical tube. To do so we overexpressed Fog in the salivary gland placode only, using *UAS-Fog x fkhGal4* (Fig. 6). Compared to the focused apical constriction near the forming invagination pit in the wild-type at early stage 11 (Fig. 6 A, A’), when Fog was overexpressed across the placode most secretory cells of the placode constricted (Fig. 6B, B’, C, C’). At late stage 11, when cells had begun to invaginate and in the wild-type a short symmetrical narrow-lumen tube has formed (Fig. 6 D,D’), in *UAS-Fog x fkhGal4* embryos the apically constricting cells had formed a large shallow depression across the whole dorsal part of the placode, with the beginning of a deeper invagination at the dorsal-posterior corner (Fig. 6 E, E’). Following on from this, at stage 12, this whole section usually formed a deepened very wide pit (Fig. 6F, pit marked in magenta in schematic in G). By stage 13 in *UAS-Fog x fkhGal4* placodes this pit often formed an extended invagination furrow on the surface of the placode, with a widened tube on the inside (Fig. 6H). This widened tube, at stage 15-16 when all cells had invaginated, displayed a significantly enlarged and flattened lumen compared to wild-type tubes (Fig. 6I-K).

**Figure 6.**
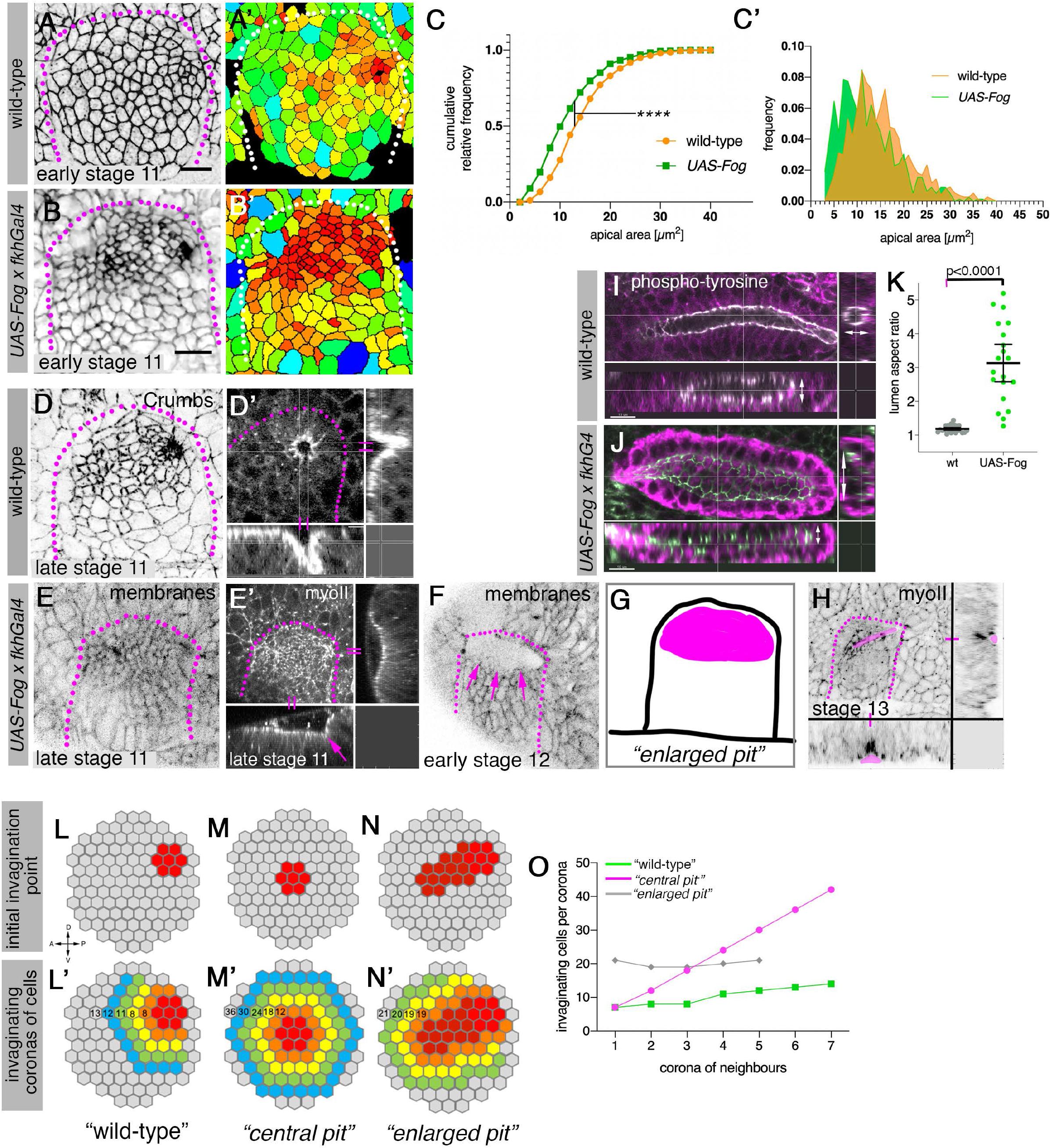
Changes to asymmetric placode patterning and invagination pit shape lead to misshapen organs. **A-C’** Whereas in the wild-type (**A**, **A’**) apical constriction commences only at the dorsalposterior corner prior to the start of pit invagination, when Fog is overexpressed (using *UAS-Fog x fkhGal4;* **B**, **B’**) excess apical constriction occurs all across the dorsal side of the placode. **A**, **B** show cell apices labelled by anti-phospho-tyrosine labelling, **A’**, **B’** show apical area size. Coloured scale for cell area as in Figure 1. **C, C’** Widening the Fog expression domain leads to significantly more constricted apices: cell area distribution of *UAS-Fog x fkhGal4* differs from wild type (Kolmogorov-Smirnov test, p-value < 0.0001). Number of cells: wild-type, n= 771 from 7 embryos; *UAS-Fog x fkhGal4*, n = 788 from 5 embryos. **D-H** In contrast to the narrow symmetrical pit that forms the early invagination in the wild-type (**D-D’**), in *UAS-Fog x fkhGal4* embryos cells in all of the whole Fog-overexpression domain constrict and initially start to form a large shallow depression (**E, E’**). **E**, **E’** are stills from a time-lapse movie, with **E’** showing surface view and xz/yz-cross sections, the arrow in **E’** points to the corner from where cells start to invaginate. The large depression is still present at stage 12 whilst cells invaginate through a large pit (**F**; magenta arrows). **G** shows a schematic of the large area (magenta) that shows concomitant apical constriction when Fog is overexpressed in the placode. The boundary of the placode and ventral midline are drawn. At stage 13, when the initial large area of constricting cells had invaginated, the pit on the surface of the embryo resembled a very extended groove (**H**; magenta area in surface view and cross sections). **I-K** At late stage 15, early stage 16, when salivary gland invagination and morphogenesis has finished, the final shape, and in particular lumen shape, of salivary gland cells overexpressing Fog (**J**; *UAS-Fog x fkhGal4)* is altered compared to wild-type controls (**I**). Apical adherens junctions are marked by phospho-tyrosine labelling (white). Glands are shown in three orthogonal cross sections. **K** Quantification of lumen aspect ratio, allowing identification of altered tube shapes such as widened tubes (Mann-Whitney test, p<000.1; wild-type: n = 23 embryos, *UAS-Fog x fkhGal4* n=20 embryos). **L-O** Theoretical considerations on how altering the shape and size of the original invagination pit will affect the geometry and shape of the invaginating and invaginated tube. **L, L’** In the wild-type, a focussed eccentric pit (red in **L**) leads to similar numbers (**L’**) of cells invaginating within each corona even in the absence of any cell rearrangements (green curve in **O**). **M, M’** A central pit would lead to an increasing number of cells invaginating in each corona, leading to a widening tube (magenta curve in **O**). **N, N’** An enlarged initial pit would lead to an increased but steady number of cells (**N’**) invaginating (grey curve in **O**), again leading to a tube with enlarged lumen.

We set out to explain the link between an enlarged or widened invagination pit in the placode early during tube formation and the expanded lumen tube observed at late stages by considering why the invagination pit forms in its eccentric position in the wild-type. We formulated a simplistic model that assumes that following the specification of an initial group of cells that apically constrict and form the initial invagination (Fig. 6L-N), neighbouring coronae of cells will follow and invaginate (Fig. 6 L’-N’). Using an idealised version of apices of a wild-type placode with an eccentric pit, the number of cells invaginating, going from corona to corona, remains very steady and low (Fig. 6L’ and green curve in O). In contrast, a centrally located pit (Fig. 6M) would lead to an ever-increasing number of cells per corona that would need to invaginate (Fig. 6M’ and magenta curve in O). An enlarged pit, a combination of the two previous scenarios (Fig. 6N) would lead to the invagination of a large number of cells initially, followed by similarly large numbers in each corona (Fig. 6N’ and grey curve in O). In fact, in these latter scenarios, the enlarged pit (Fig. 6N, N’) resembles aspects of the enlarged pit and invagination seen in *UAS-Fog x fkhGal4* embryos. Furthermore, the central pit resembles the events observed in some *hkb^-/-^* mutant embryos, where, as shown above, a delayed central area of constriction forms and turns to a widened invagination (Suppl. Fig. 3_2). This central and enlarged pit at late stages often collapsed into an enlarged crescent of subduction at the boundary (Supplemental Fig. 3_2 and Supplemental Movies 5 and 6), leading to malformed and enlarged lumen glands at the end of embryogenesis, as has been observed previously (9).

Thus, the eccentrically placed invagination pit in the dorsal-posterior corner of the salivary gland placode is a key factor in ensuring the formation of a symmetrical narrow lumen tube during invagination, in part likely due to the control over the number of invaginating cells in each corona destined to internalise.

## Discussion

In this work, we investigated how the sustained morphogenesis of a tissue is achieved, in this case the continued invagination of epithelial cells forming a tubular organ. We previously identified cell behaviours that become established across the placode and drive the initial tissue bending (6).

A number of mechanisms appear at play that in combination ensure that a symmetrical and narrow-lumen tube is formed from an initially flat epithelial primordium. A temporal and spatial component is provided by the dynamically changing pattern of the transcription factors Hkb and Fkh. These are expressed initially with the highest protein level at the point where the invagination pit will form, before expression expands further across the placode over time. This ensures that the cells in the dorsal-posterior corner are the first to experience high levels of activity of these transcription factors, and we show that one of the important targets, the GPCR-ligand Fog, is found at high levels in these cells in a Hkb- and Fkh-dependent manner with a slight temporal delay. A further spatial component is provided by the fact that, as cells continuously internalise to form the tube on the inside of the embryo, new coronas of cells are brought into a ‘near the pit’ position and ‘adjust their behaviour’ accordingly. This position-specific behaviour of apical constriction is most likely triggered by the previous temporal expansion of Hkb and Fkh expression to these cells, thereby priming them for their next step in the path to tube formation. Interestingly, though, the pattern of Fkh and Hkb expression from t = 0 min onwards, though generally expanding across the placode, is not identical. This could suggest that downstream targets begin to diverge at later stages.

Our work strongly suggests that the combined early patterning of Hkb and Fkh expression defines the early invagination pit and its eccentric position. How though is Hkb and Fkh expression initially limited to this point? Both are expressed in the placode downstream of the upstream homeotic factor Sex Combs Reduced (Scr; (8, 18)). They are also not the only factors expressed near exclusively at this corner first. At the transcription factor level in particular one of the parasegmentally repeated stripes of Wingless (Wg) expression overlays the posterior part of the placode, its expression domain defined by previous parasegmental patterning (19). And at the time point when Hkb and Fkh expression at the future pit location commences, the Wg expression within the placode region is also confined to this corner only, rather than extending all the way to the ventral midline (data not shown). Future studies will show whether a combination of Scr and Wg transcription factor activity could activate early pit-defining expression of factors such as Fkh and Hkb. In addition to transcription factors, some potential downstream effectors also appear to be expressed near the future pit location first, these include Btk29/Tec29 (11), Crumbs (9, 20), 18-wheeler (21), and Klarsicht (9). Cell-based genomics approaches such as single cell-RNAseq or ATAC-seq should in the future allow to determine precisely the genetic programme that underlies the regionalisation of the placode.

Activation of actomyosin contractility through GPCR signalling driven by the autocrine secretion of the ligand Fog is a module repeatedly used during tissue bending and invagination events in *Drosophila* embryogenesis. What differs are the upstream activating transcription factors: during mesoderm invagination Fog expression on the ventral side is controlled by Twist and Snail (22), whereas its expression during posterior midgut invagination is controlled by Tailless and Hkb (23). The effect of Hkb on Fog expression is likely direct, as the Fog locus contains predicted Hkb-binding sites (using oPOSSUM, http://opossum.cisreg.ca/oPOSSUM3/). The identity of the GPCR that could mediate the Fog signal in the salivary gland placode is unclear thus far. Published and database (Flybase) *in situs* for the GPCRs previously reported to be involved in mesoderm invagination and germband extension, Mist and Smog (24, 25), do not show an enriched expression in the placode. But with around 200 *Drosophila* GPCRs in existence, half of them orphan receptors (26), the placode might well express yet another one, and we will focus on candidates previously identified in screens (27) and genomics approaches (our unpublished data) in the future.

The question of how cell behaviour is modulated over time after the initial transcriptional patterning that defines primordium identity is key to understand general mechanisms that allow continuous organ formation. Different solutions seem to have been adopted in different tissues. Recent work analysing mesoderm invagination in *Drosophila* has shown that the key morphogenetic effectors T48 and Fog show a gradient of expression due to differential timing of transcriptional activation, leading to different spatial accumulations of the transcripts (28) and proteins (29). Flattening the gradient experimentally (by expansion of the ventral domain) leads to a widened and perturbed invagination process. By contrast, the invagination of the posterior midgut or endoderm primordium in the fly embryo does not require sustained transcription, but instead a wave of apical constriction that is initiated by Fog expression in a subset of cells, appears to be propagated via mechanical feedback (30). The salivary gland placode seems to follow a model of a temporal expansion or wave of transcriptional activity translating into positionally fixed behaviours. Nonetheless, our quantitative analysis of 3D cell behaviours strongly indicated that mechanical feedback does also play a role (6). Which combination of mechanisms is ultimately at work to sustain a morphogenetic process could depend on a number of factors including the developmental stage, the speed of the process (consider ~15 min for mesoderm invagination versus ~30 min for endoderm invagination versus ~1.5 hours for secretory cell invagination in the salivary gland placode), as well as the influence of other nearby concomitant morphogenetic events.

Curiously and maybe tellingly, the asymmetry of the placodal set-up is retained in another tube formation process, the invagination of the posterior spiracles (31, 32). The posterior spiracle placode, like the salivary gland placode, is a flat epithelium with an eccentric invagination point. In this case an initial tube is already present due to the previous invagination of the most posterior tracheal placode. Whether the asymmetrical set-up is thus a conserved feature of further tube budding processes during development will be an exciting question to address. Mathematical modelling of tube morphogenesis assuming an eccentric versus centrally positioned invagination point will help to dissect the topological advantages of being asymmetric.

In summary, we have quantitatively unravelled a dynamic spatio-temporal patterning of transcription factors and switches in cell behaviours leading to positionally fixed behaviours during the morphogenesis of simple tubular organ. We foresee that such mechanisms will be important for establishing general mechanisms and morphogenetic modules at work during the morphogenesis of more complex tubular organs and potentially also for engineered tissues in a dish.

## Acknowledgments

The authors would like to thank the following people; for reagents and fly stocks: Debbie Andrew, Thomas Lecuit, Stefan Luschnig, Herbert Jäckle, Naoyuki Fuse. We thank members of the lab for input on the manuscript.

K.R. and Y.S.C. were supported by the Medical Research Council (file reference number U105178780). G.B. was supported by a Wellcome Trust Investigator Award (207553/Z/17/Z) to Bénédicte Sanson, Dept. of PDN, Cambridge, UK.

## Author contribution

Conceptualisation, K.R. and Y.S-C..; Methodology, K.R., Y.S-C. G.B.; Investigation, K.R., Y.S-C.; Writing-Original Draft, K.R., Y.S-C. Funding Acquisition, K.R.

## Materials & Methods

### Key Resources Table

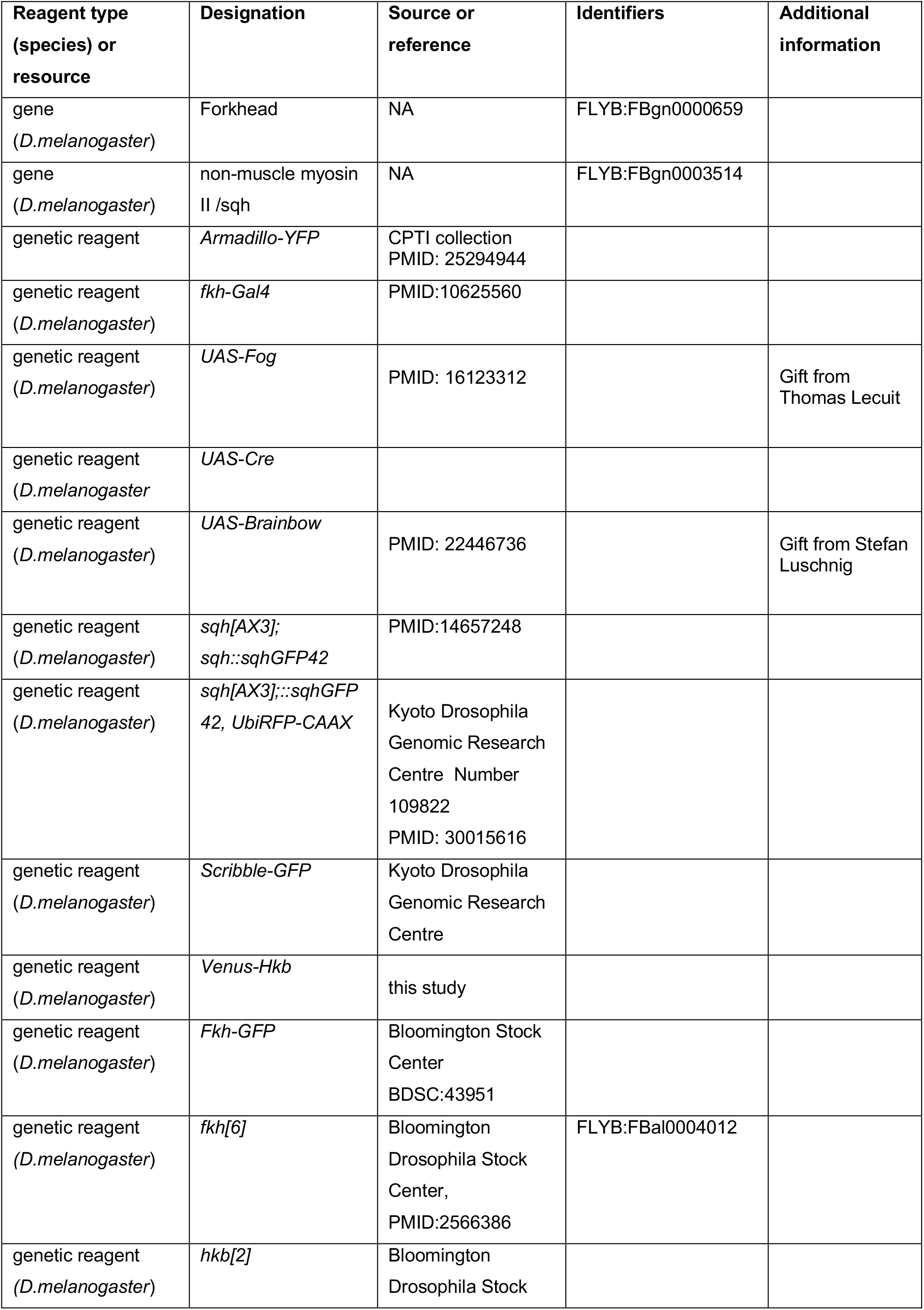
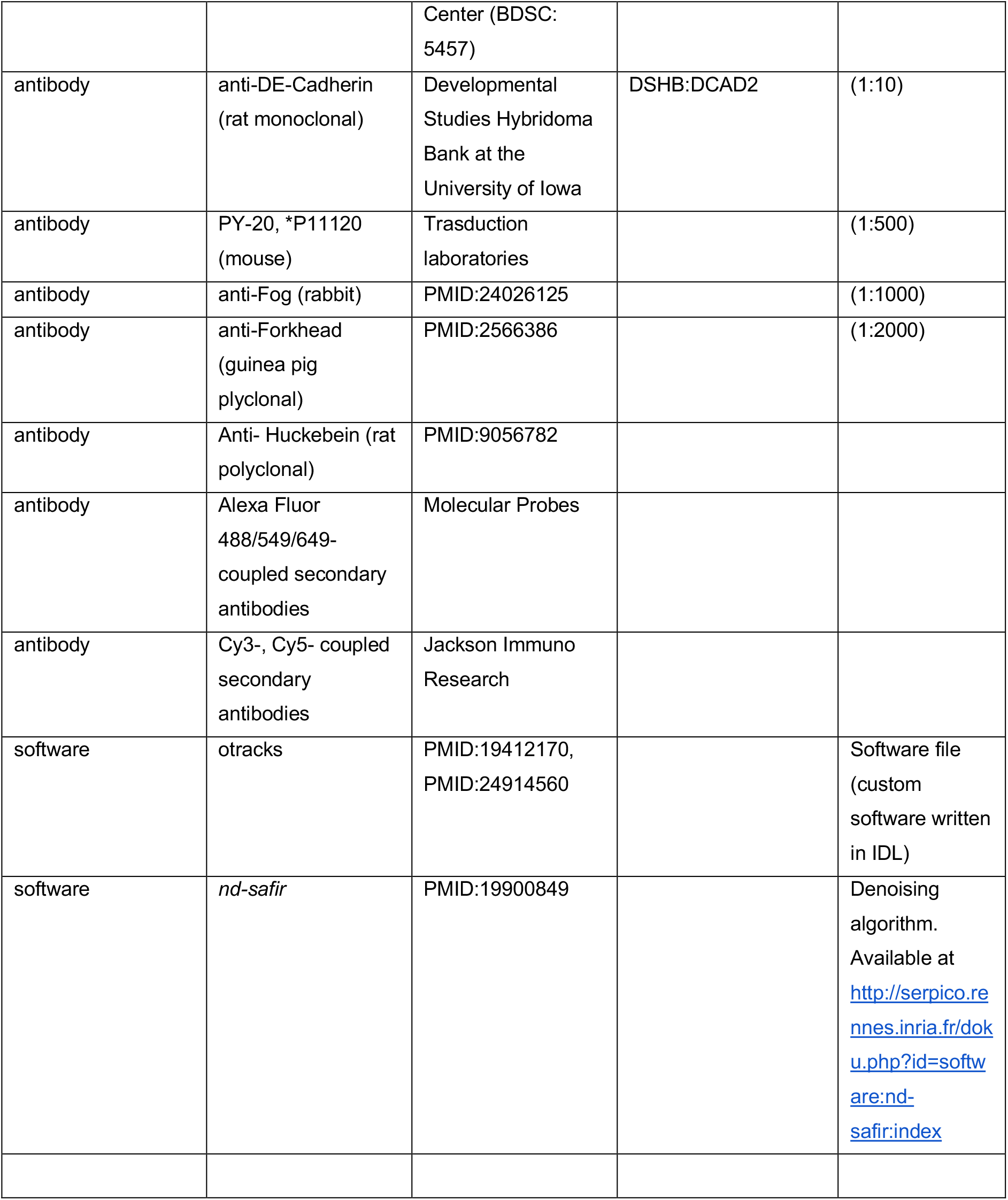

### Fly stocks and husbandry

The following transgenic fly lines were used: *Armadillo-YFP* (CPTI collection described in (33)), *sqhAX3; sqh::sqhGFP42* (34) and *fkhGal4* (35, 36) [kind gift of Debbie Andrew]; *UAS-Fog* (kind gift of Thomas Lecuit); *y*^1^ *w*\* *cv*^1^ *sqh^AX3^; P{w^+mC^=sqh-GFP.RLC}C-42 M{w^+mC^=Ubi-TagRFP-T-CAAX}ZH-22A* (Kyoto DGRC Number 109822, referred to as *sqh^AX3^;sqhGFP; UbiRFP); [sqhGFP42,UbiRFP; fkh[6]/TM3 Sb Twi-Gal4::UAS-GFP] (fkh[6]* allele from Bloomington *J;[hkb^2^, p^2^, Scribble-GFP]; [sqhGFP42,UbiRFP; hkb^2^, p^2^ / TM3 Sb Twi-Gal4::UAS-GFP] hkb^2^* allele from Bloomington BDSC: 5457), [*UbiRFP, Venus-HKB]* (this study), *FkhGFP* (Bloomington BDSC:43951) and *[FkhGFP; wgGal4, UAS-palmYFP]* (generated from membrane Brainbow (37); (38)). See Table 1 for details of genotypes used for individual figure panels.

**Table 1.**
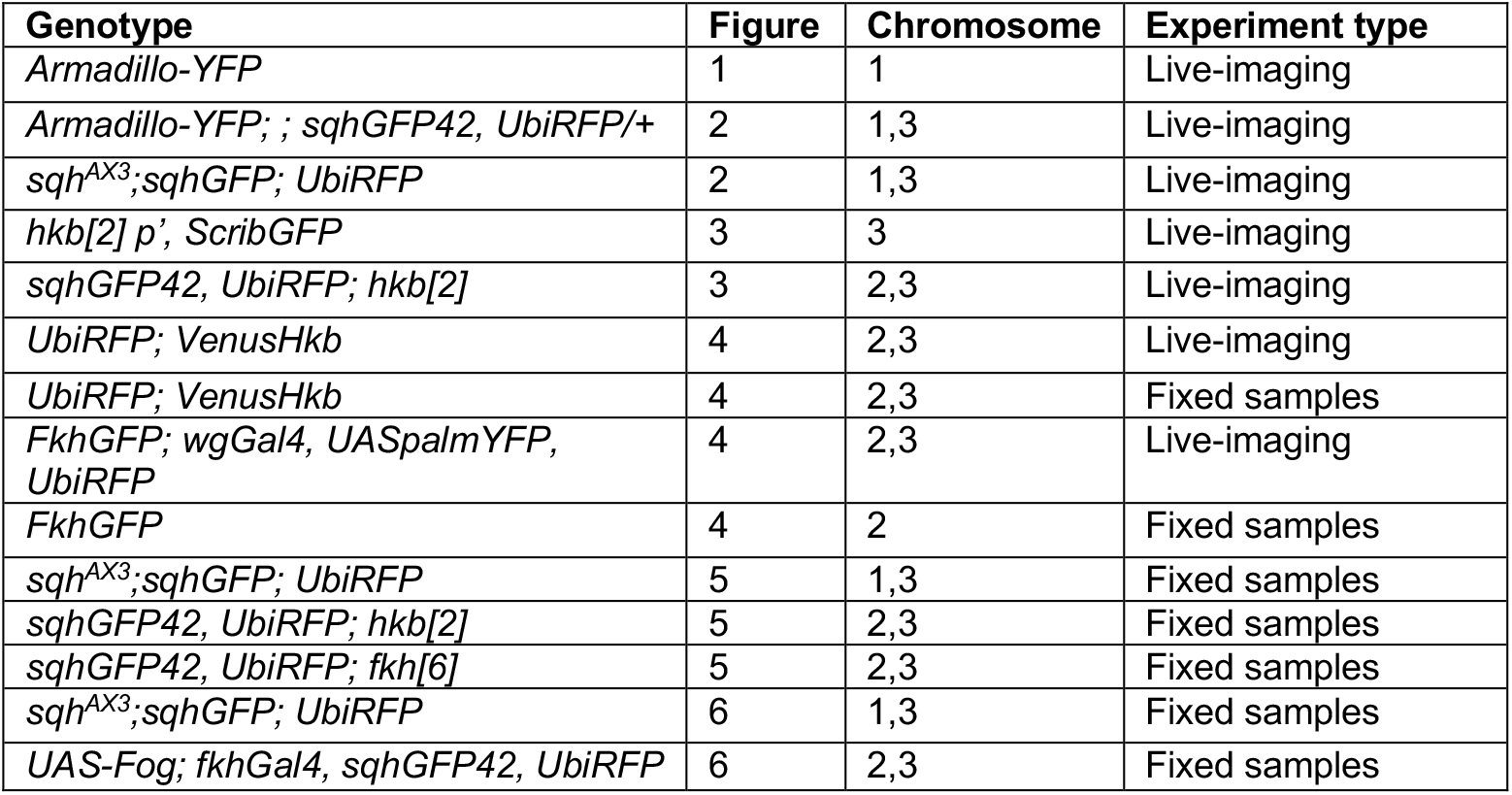
Embryo genotypes as presented in figures.

### Embryo Immunofluorescence Labelling, Confocal, and Time-lapse

Embryos were collected on apple juice-agar plates and processed for immunofluorescence using standard procedures. Briefly, embryos were dechorionated in 50% bleach, fixed in 4% formaldehyde, and stained with primary and secondary antibodies in PBT (PBS plus 0.5% bovine serum albumin and 0.3% Triton X-100). Anti-Hkb was a gift from Chris Doe (39); anti-Fkh was a gift from Herbert Jäckle (40) and anti-Fog was a gift from Naoyuki Fuse (41), PY- 20, (*P11120, Transduction laboratories); anti-CrebA (DSHB). Secondary antibodies used were Alexa Fluor 488/Fluor 549/Fluor 649 coupled (Molecular Probes) and Cy3 coupled (Jackson ImmunoResearch Laboratories). Samples were embedded in Vectashield (Vectorlabs).

Images of fixed samples were acquired on an Olympus FluoView 1200 or a Zeiss 780 Confocal Laser scanning system as z-stacks to cover the whole apical surface of cells in the placode. Z-stack projections were assembled in ImageJ or Imaris (Bitplane), 3D rendering was performed in Imaris.

For live time-lapse experiments (see Table 1), embryos were dechorionated in 50% bleach and extensively rinsed in water. Stage 10 embryos were manually aligned and attached to heptane-glue coated coverslips and mounted on custom-made metal slides; embryos were covered using halocarbon oil 27 (Sigma) and viability after imaging after 24h was controlled prior to further data analysis. Time-lapse sequences were imaged under a 40x/1.3NA oil objective on an inverted Zeiss 780 Laser scanning system, acquiring z-stacks every 0.58-3 minutes with a typical voxel xyz size of 0.22 x 0.22 x 1 μm. Z-stack projections to generate movies in Supplementary Material were assembled in ImageJ or Imaris. The absence of fluorescent *Twi-Gal4::UAS-GFP* was used to identify homozygous *hkb[2]* and *fkh[6]* mutant embryos. The membrane channel images from time-lapse experiments were denoised using *nd-safir* software (42).

### Cell segmentation and tracking

Cell segmentation and tracking was performed using custom software written in (13). First, the curved surface of the embryonic epithelium was located by draping a ‘blanket’ over all image volumes over time, where the pixel-detailed blanket was caught by, and remained on top of binarised cortical fluorescence signal. Quasi-2D image layers were then extracted from image volumes at specified depths from the surface blanket. We took maximum intensity projections of a small number of near surface image layers typically at 0-4 μm from membrane channels, for cell-apical analyses, apicomedial Myosin and anti-Fog fluorescence channels and projections of image layers typically from 5-16μm encompassing nuclear fluorescence for Venus-HKB and FkhGFP analyses. Membrane channels were filtered with median, top-hat or high/low frequency filters as necessary to optimise subsequent cell tracking.

Cells in membrane channels were segmented using an adaptive watershedding algorithm as they were simultaneously linked in time. Manual correction of segmented cell outlines was performed for all fixed and time-lapse data. The segmentation of all the movies used in this study was manually corrected to ensure at least 90% tracking coverage of the placode at all times. Tracked cells were subjected to various quality filters (lineage length, area, aspect ratio, relative velocity) so that incorrectly tracked cells were eliminated prior to further analysis. The number of embryos analysed and number of cells can be found in Supplemental Figures 1 and 3.

### Mobile radial coordinate system for the salivary gland placode

Wild-type movies were aligned in time using as t= 0min the frame just before the first sign of invagination at the future tube pit was evident. *hkb^-/-^* mutants were aligned using as a reference of embryo development the level of invagination of the tracheal pits that are not affected in the *hkb^-/-^* mutant as well as other morphological markers such as appearance and depth of segmental grooves in the embryo. Cells belonging to the salivary placode (without the future duct cells that comprise the two most ventral rows of cells in the primordium) were then manually outlined at t= 0 min using the surrounding myosin II cable as a guide and ramified forwards and backwards in time. Only cells of the salivary gland placode were included in subsequent analyses.

At t= 0 min, the centre of the future tube pit was specified manually as the origin of a radial coordinate system, with radial distance (in μm) increasing away from the pit. Circumferential angle was set to zero towards Posterior, proceeding anti-clockwise for the placode on the left-hand side of the embryo, and clockwise for the placode on the right so that data collected from different sides could be overlaid.

The radial coordinate system was ‘mobile’, in the sense that its origin tracked the centre of the pit, forwards and backwards in time, as the placode translated within the field of view due to embryo movement or to on-going morphogenesis.

### Morphogenetic strain rate analysis

Detailed spatial patterns of the rates of deformation across the placode and over time quantify the net outcome of active stresses, viscoelastic material properties and frictions both from within and outside the placode. We quantified strain (deformation) rates over small spatio-temporal domains composed of a focal cell and one corona of immediate neighbours over a ~5 min interval ((13) and reviewed in (43)). On such 2D domains, strain rates are captured elliptically, as the strain rate in the orientation of greatest absolute strain rate, with a second strain rate perpendicular to this.

For the early morphogenesis of the salivary gland placode, in which there is no cell division or gain/loss of cells from the epithelium, three types of strain rate can be calculated. First, total tissue strain rates are calculated for all local domains using the relative movements of cell centroids, extracted from automated cell tracking. This captures the net effect of cell shape changes and cell rearrangements within the tissue, but these can also be separated out. Second, domain cell shape strain rates are calculated by approximating each cell with its best-fit ellipse and then finding the best mapping of a cell’s elliptical shape to its shape in the subsequent time point, and averaging over the cells of the domain. Third, intercalation strain rates that capture the continuous process of cells in a domain sliding past each other in a particular orientation, is calculated as the difference between the total tissue strain rates and the cell shape strain rates of cells. Strain rates were calculated using custom software written in in L3Harris Geospatial IDL (code provided in (13) or by email from G.B.B.).

The three types of elliptical strain rate were projected onto our radial coordinate system (6), so that we could analyse radial and circumferential contributions. Strain rates in units of proportional size change per minute can easily be averaged across space or accumulated over time. We present instantaneous strain rates over time for spatial subsets of cells in the placode (Supplemetal Figure 3_3), and cumulative strain ratios for the same regions over time. These plots were made from exported data using MATLAB R2014b. To test for differences in instantaneous strain rates of wt and *hkb^-/-^* time-lapse movies we used a mixed-effects model implemented in R (lmer4 package as in (4, 6, 44)) with a significance threshold of p<0.05. The phenotype (wt or *hkb^-/-^*) was considered a fixed effect while the variation between embryos from the same phenotype was considered a random effect.

### Neighbour exchange analysis

We used changes in neighbour connectivity in our tracked cell data to identify neighbour exchange events (T1 processes). Neighbour exchange events were defined by the identity of the pair of cells that lost connectivity in *t* and the pair that gained connectivity at *t+1*. The orientation of gain we defined as the orientation of the centroid-centroid line of the gaining pair at *t+1*. We further classified gains as either radially or circumferentially oriented, depending on which the gain axis was most closely aligned to locally. We did not distinguish between solitary T1s and T1s involved in rosette-like structures.

From visual inspection, we knew that some T1s were subsequently reversed, so we characterised not only the total number of gains in each orientation but also the net gain in the circumferential axis, by subtracting the number of radial gains. Furthermore, when comparing embryos and genotypes, we controlled for differences in numbers of tracked cells by expressing the net circumferential gain per time step as a proportion of half of the total number of tracked cell-cell interfaces in that time step. We accumulated numbers of gains, net gains, and proportional rate of gain over time for WT and *hkb* embryos (Figure 3L,M). Two sample Kolmogorov-Smirnov tests were used to determine significance at p<0.05 for data in Figure 3.

### Automated medial myosin II quantification and oscillatory behavior

We extracted a quasi-2D layer image from the myosin II channel at a depth typically from 0-3 μm that maximised the capture of apical-medial myosin. We background-subtracted the myosin images and quantified the average intensity of apical-medial myosin as the fluorescence inside the segmented cell, excluding the fluorescence along cell-cell interfaces encompassing the cell edge pixels plus 2 pixels in a perpendicular direction either side (considered junctional myosin).

The analysis of apical-medial fluorescence fluctuations was performed as described before in (45) and (7). Briefly, the apical-medial fluorescence values in the time series were detrended by subtracting a boxcar average with a window of 6 minutes (larger than the maximum expected fluctuation cycle length). All peaks and troughs were then identified in the fluorescence signal. Peaks or troughs that were associated with very small amplitude cycles were skipped. The remaining peaks and troughs were used to calculate the amplitude and the cycle length (peak to peak) and the amplitude (average peak to trough) of fluctuations.

### Hkb, Fkh and Fog fluorescence intensity quantifications

Average fluorescence was calculated after background subtraction as the mean fluorescence inside a segmented cell excluding the cell edge pixels and 2 pixels in a perpendicular direction either side. Fixed samples of *Venus-Hkb* and *FkhGFP* embryos of mid to late stage 10 (t<< 0min), early stage 11 (t+ 0min) and late stage 11 (t>> 0 min) were pooled together with relevant time-points from live-imaging as shown in Table 1 and figure legends. Fluorescence was normalised in each image by dividing the average fluorescence of each cell by the 98^th^ percentile value. Calculations were performed in R.

---

**Supplemental Figure S1, related to Figure 1.**
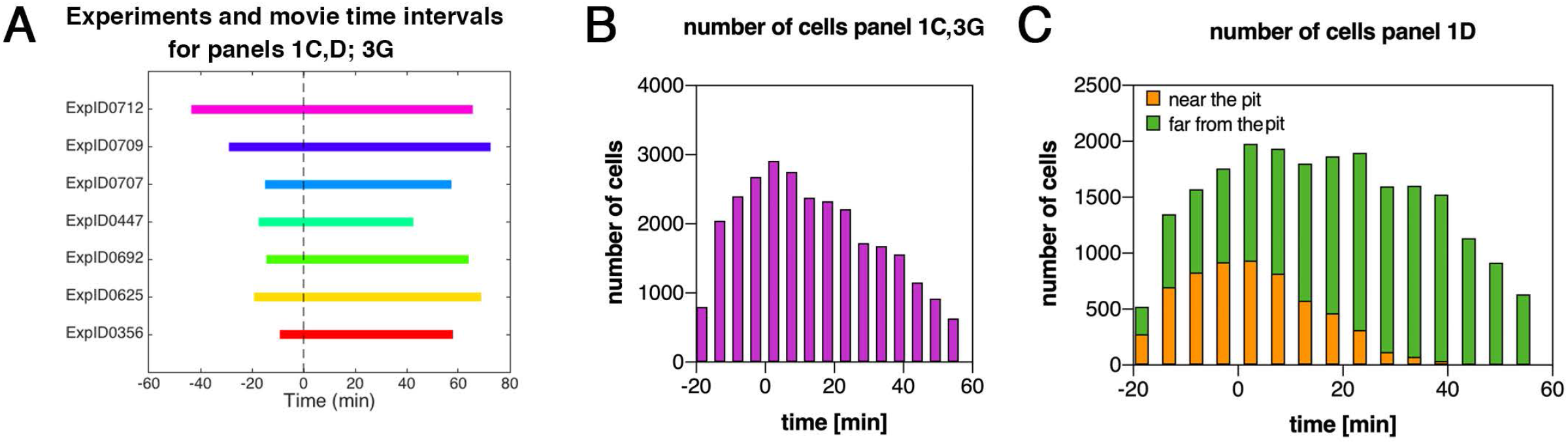
Patterned apical constriction remains fixed around the pit over time. **A** Illustration of time intervals covered by wild-type time-lapse movies analysed in Fig. 1C and D, and also in Fig. 3G. **B** Number of cells used in wild-type experiments per time interval to calculate cell area change shown in Fig. 1C and 3G. **C** Numbers of cells analysed over time, split into near the pit and far from the pit located cells, analysed for Fig. 1D.

**Supplemental Figure S2, related to Figure 2.**
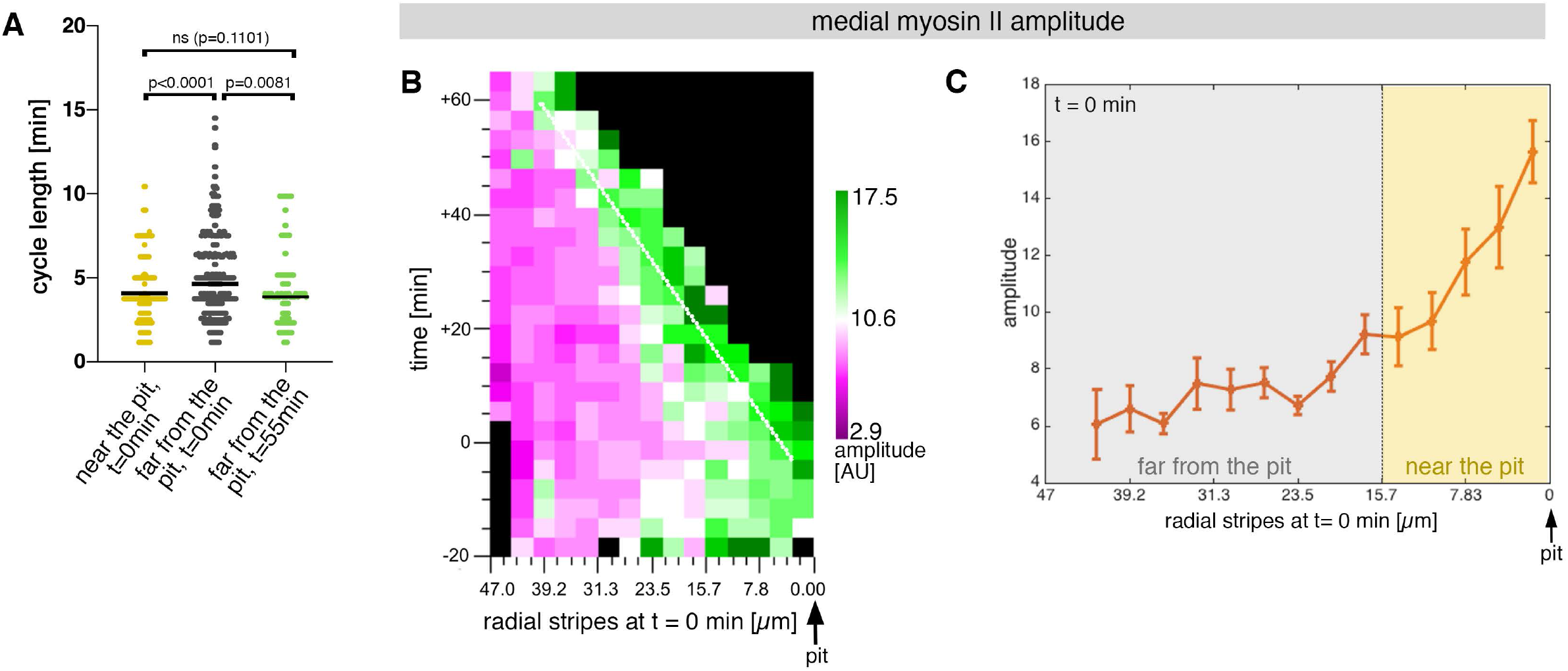
Sustained apical constriction near the pit is driven by apical-medial myosin. **A** Myosin cycle length for all cells analysed from 3 movies, split into cells located near the pit (n=148) compared to cell located far form the pit (n=302) at t=0. Cells near the pit have significantly higher shorter, myosin cycle length than cell located far from the pit; statistical significance as determined using Mann Whitney test, with p<0.0001. The myosin cycle length of cells far from the pit at later time points (t=55-60 min, n = 134) was not significantly different from cells near the pit at t=0 (p = 0.1101 using a Mann Whitney test). Related to Figure 2. **B** Spatial representation of the distribution of the myosin amplitude across the salivary gland placode, with radial location (collapsed into stripes) of cells specified at t= 0min. The dashed line marks the region of highest amplitude that is always adjacent to the invagination pit as cells flow into it over time. The mean of 3 movies is shown. **C** Spatial representation of the amplitude of myosin oscillations in radial stripes from the pit (0 μm) to the boundary of the placode (47μm) at t= 0min. Mean and standard error of the mean are shown. Data are pooled from 3 movies ranging from −7.5-56.25 min; −16.76 to 55.48 min and −17.42 to 67.26min. Regions corresponding to cells near the pit and far from the pit are indicated by coloured shading.

**Supplemental Figure S3_1, related to Figure 3.**
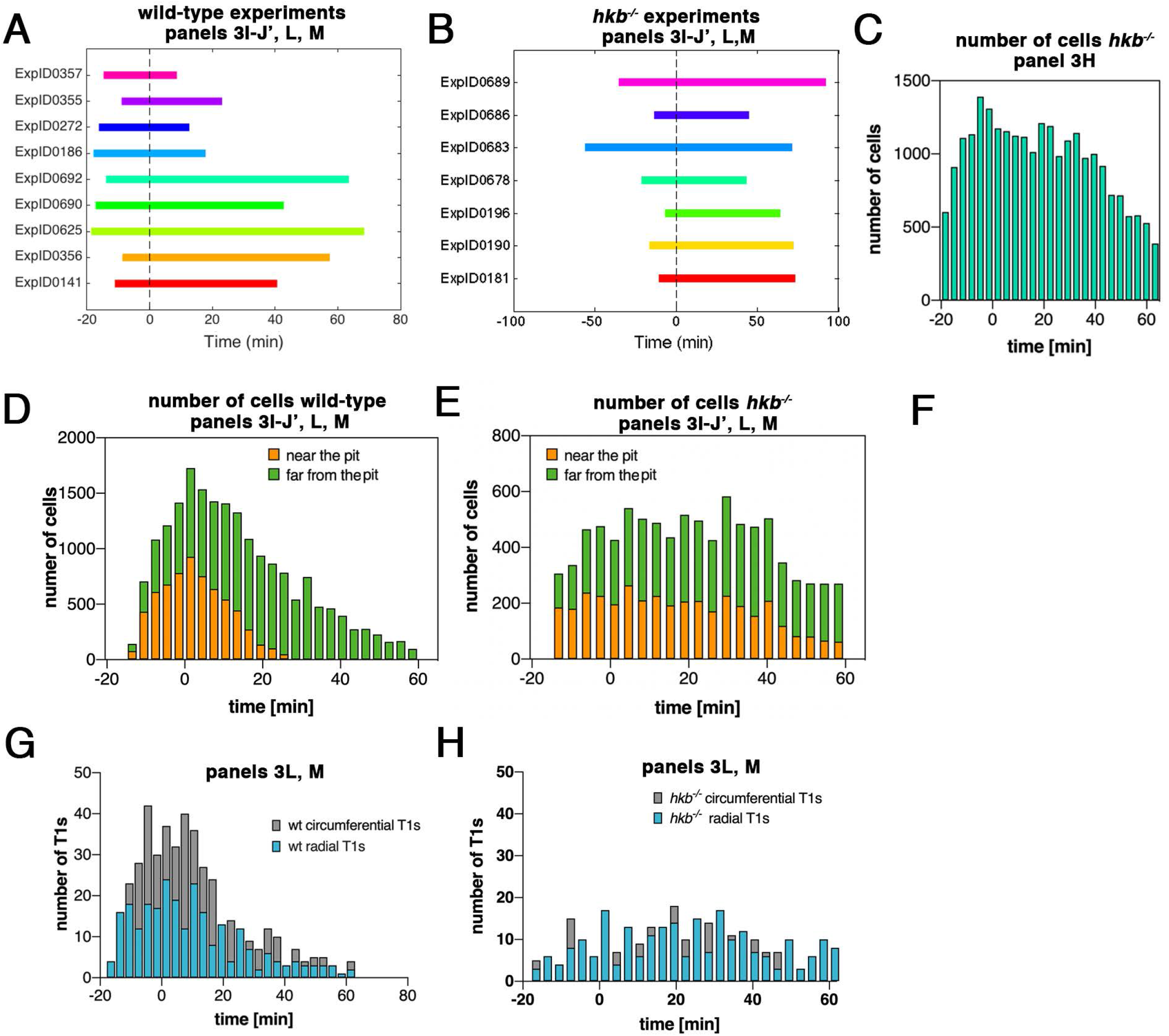
hkb^-/-^ mutants show a delayed symmetrical apical constriction. **A** Illustration of time intervals covered by the time-lapse movies for wild-type experiments used to calculate strain rates in Fig. 3I-J’ and T1 productive neighbour exchanges in Fig. 3L, M. For cell area change in the wild-type shown in Fig. 3G, the experiments ExpID356, ExpID0625, ExpID0692, ExpID0447, ExpID0707, ExpID0709, ExpID0712 were analysed (details of time interval for the last 4 experiments are shown in Suppl. Fig. 1). **B** Illustration of time intervals covered by the time-lapse movies for *hkb^-/-^* mutant experiments. All experiments shown were used to calculate the cell area change shown in Fig. 3H. Experiments ExpID0181, ExpID0190, ExpID0196, ExpID0689 and ExpID0678 were used to calculate strain rates in Fig. 3I-J’ and T1 neighbour exchanges in Fig. 3L, M. **C** Number of cells used in *hkb^-/-^* mutant experiments per time interval to calculate cell area change shown in Fig, 3H. **D, E** Number of cells in wild-type experiments (**D**) and *hkb^-/-^* mutant experiments (**E**) used to calculate strain rates in Fig. 3I-J’ and T1 exchanges in Fig. 3L, M by time interval and region (near the pit and far from the pit). **G, H** Total number of T1 exchanges detected in wild-type experiments (**G**) and *hkb^-/-^* mutant experiments (**H**), divided into circumferential and radial T1s. The cumulative plots of T1 exchanges are shown in Fig. 3L,M.

**Supplemental Figure S3_2, related to Figure 3.**
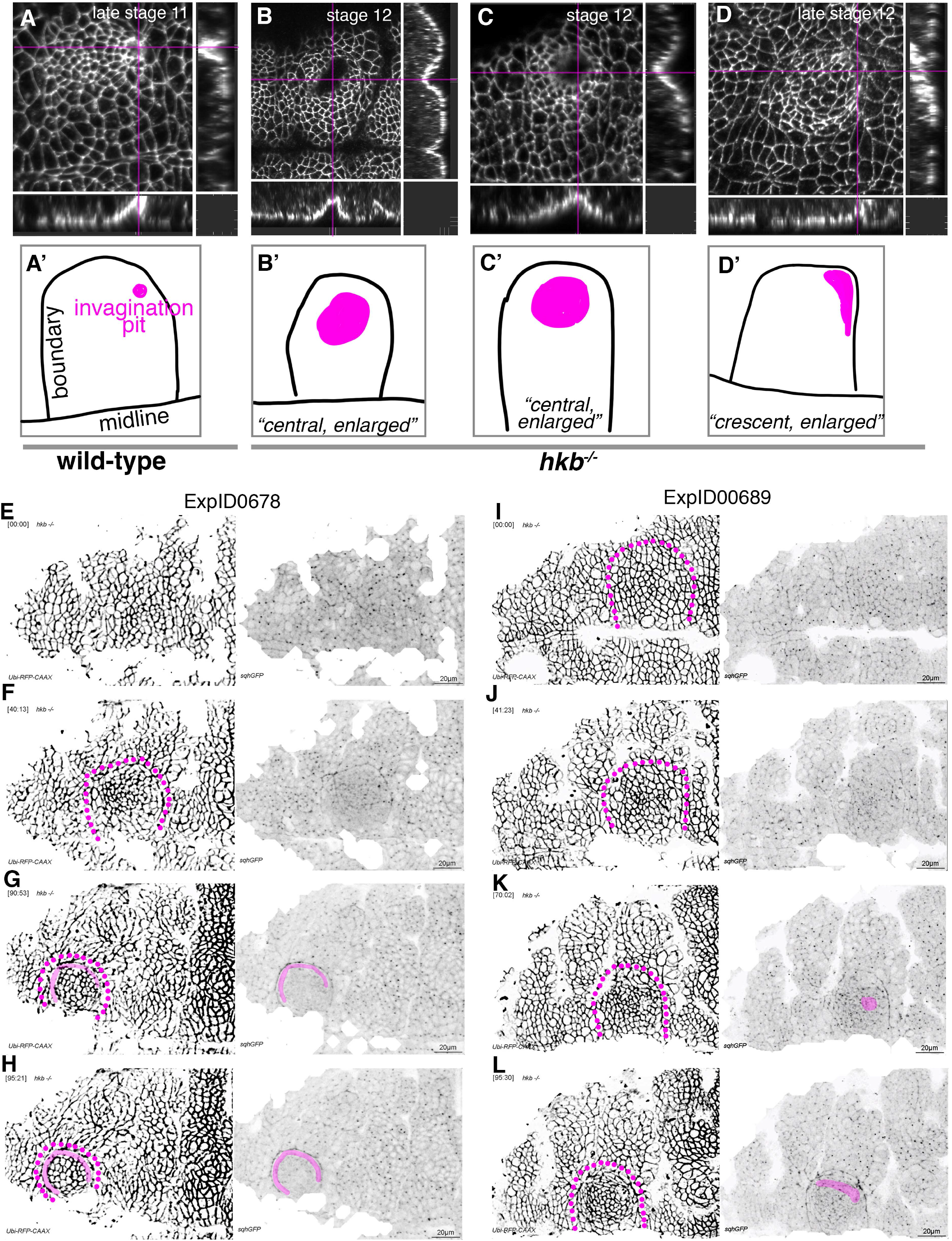
hkb^-/-^ mutants show a delayed symmetrical apical constriction followed by aberrant invagination. **A, A’** The invagination pit in wild-type embryos at stage 11 and beyond is small, eccentric in its position in the placode (dorsal-posterior corner). **B-D’** In *hkb^-/-^* mutant embryos, the invagination pit that forms at a delayed stage is often centrally positioned within the placode and enlarged, leading to a very much widened invagination as visible in the cross-sections. These central pits often enlarge towards the boundary and sometimes collapse into a more crescent-shaped invagination at later stages. Magenta crosshairs in **A-D** indicate the positions of the cross-sections shown. Labelling in these panels is for anti-phospho-tyrosine. In **A’-D’** the position and shape of the invagination pit, drawn from the images in **A-D**, is shown in magenta, and the boundary of the placode as well as the ventral midline, where visible, are in black. **E-H** Still images from time lapse movie ExpID0678 shown in Supplemental Movie 5. The delayed central constriction is visible in **F**, the crescent-shaped enlarged invagination front is highlighted in magenta in **G**, **H**. **I-L** Still images from time lapse movie ExpID0689 shown in Supplemental Movie 6. The delayed central constriction is visible in **J**, the enlarged invagination is highlighted in magenta in **K, L**. Membrane channel and myosin II channel are shown as individual panels in **E-L**, placode boundaries are indicated by magenta dotted lines.

**Supplemental Figure S3_3, related to Figure 3.**
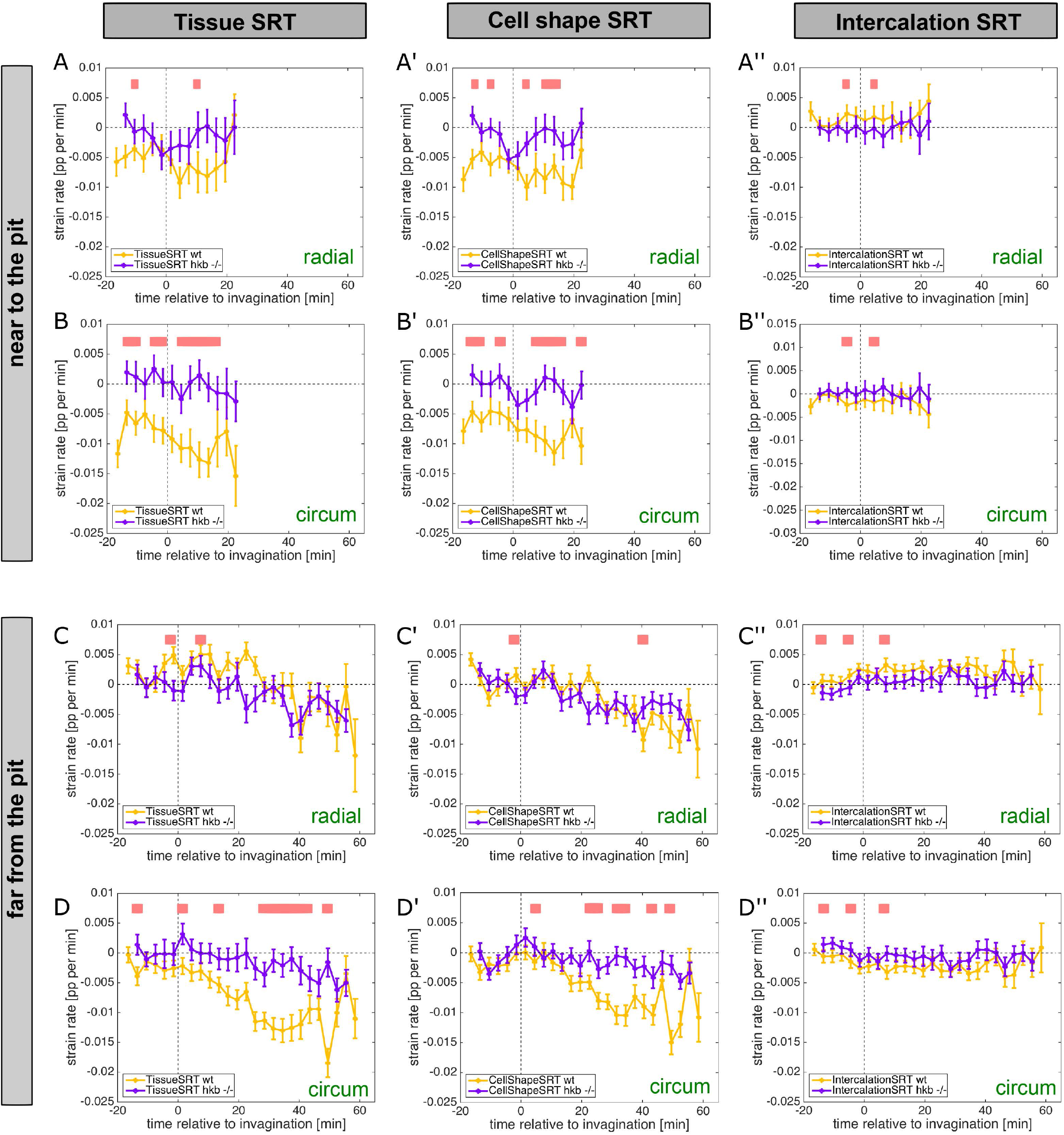
hkb^-/-^ mutants show aberrant cell behaviours. **A-D’’** Instantaneous strain rates relating to the cumulative strain rates shown in Figure 3 I-J’. Panels for cell near the pit and far from the pit are shown, split into tissue, cell shape and intercalation strain rate tensors, as well as the radial and circumferential contributions. Regions of statistically significant differences (p<0.05) are indicated by pink bars above the curves, and significance was determined using mixed-effects model.

**Supplemental Figure S4, related to Figure 4.**
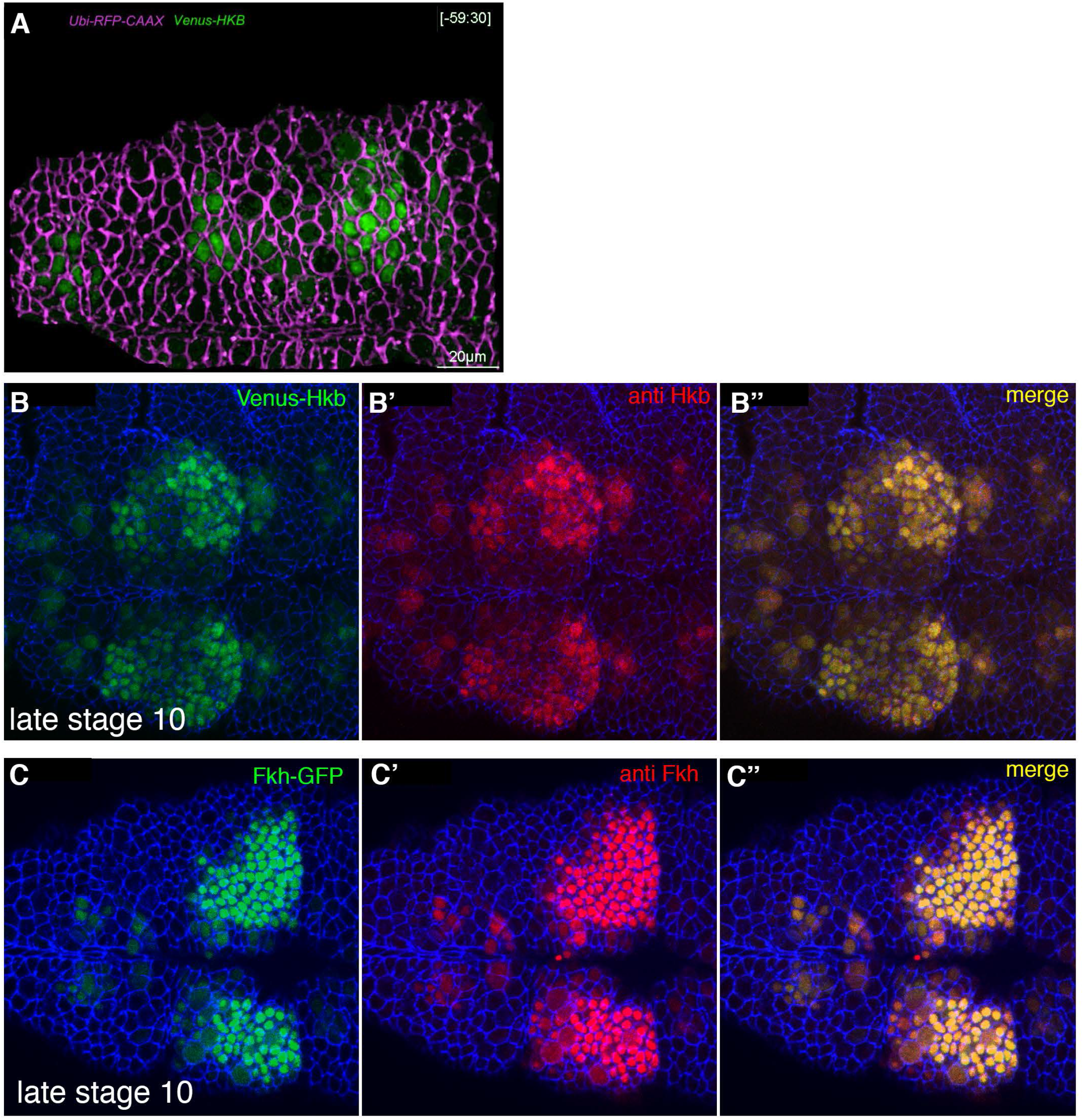
The salivary gland placodal primordium is asymmetrically patterned by Hkb and Fkh prior to morphogenesis. **A** Still of Supplemental Movie 7 at t= −59.5min. Cell membranes are shown in magenta, Venus-Hkb is in green. **B-B’** Comparison of Venus-Hkb (green in **B**) and anti-Hkb antibody staining (red in **B’**) at late stage 10. **B-B’** Comparison of Fkh-GFP (green in **B**) and anti-Fkh antibody staining (red in **B’**) at late stage 10.

**Supplemental Figure S5, related to Figure 5.**
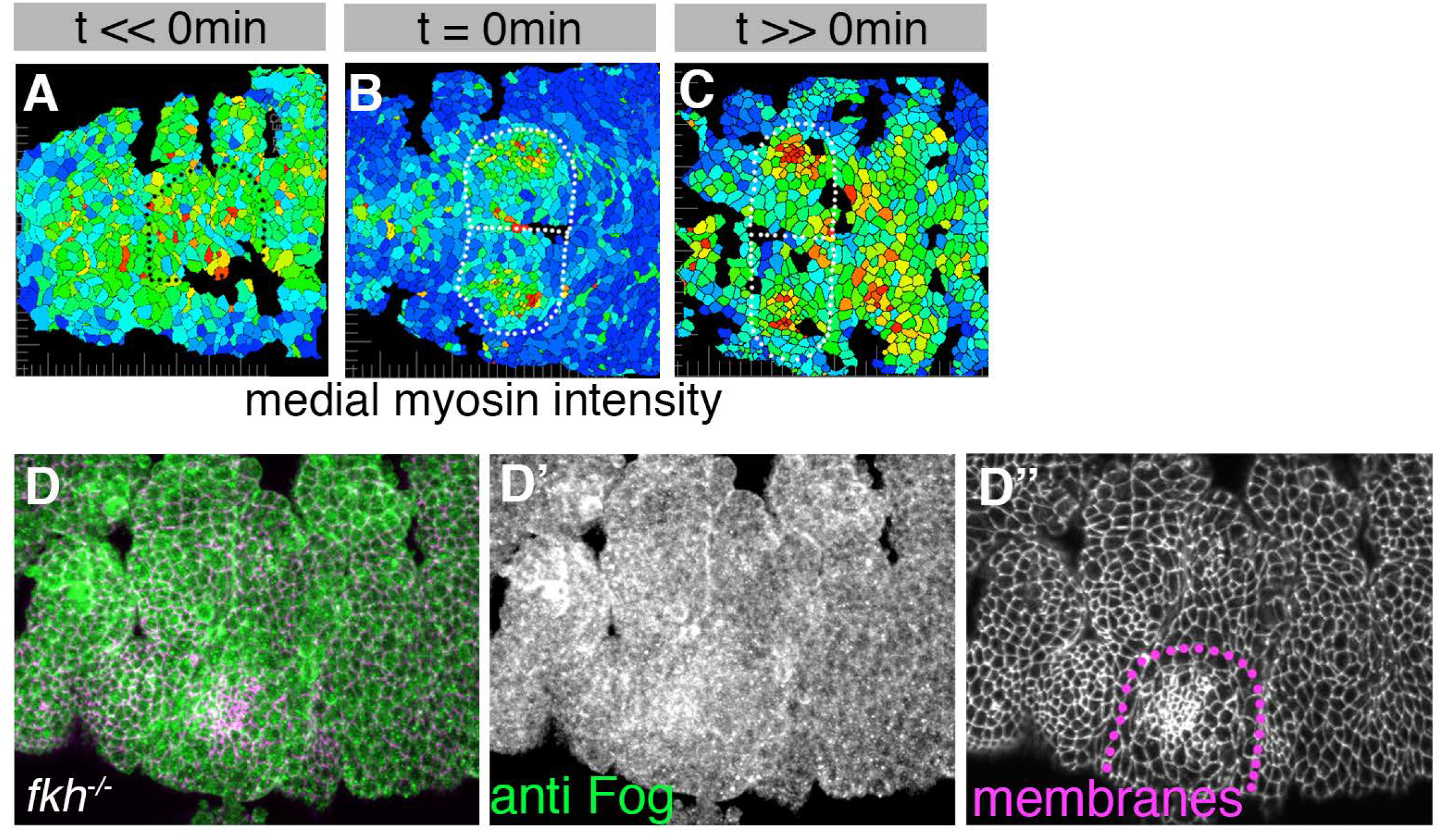
Asymmetric Fog expression is controlled by Hkb and Fkh and is upstream of early differential behaviours in the placode. **A-C** Wild-type embryos expressing sqhGFP were analysed before, at initiation of and well beyond start of pit invagination and medial myosin II intensity quantified per cell. The boundary of the placode(s) is indicated by dotted lines. Colour scale for apical cell area as in Figure 1. **D-D’’** anti-Fog antibody labeling (green in **D** and **D’**) in a *fkh^-/-^* mutant embryo, showing no increased Fog accumulation in the placode over the surrounding epidermis. Cell membranes are in magenta in **D** and in **D’’**

## List of Supplemental Movies

**Supplemental Movie 1, related to Figure 1. Contiuous apical constriction near the pit.** Time -lapse movie of an embryo expressing Armadillo-YFP to label cell outlines. Apical cell area change is colour-coded.

**Supplemental Movie 2, related to Figure 2. Continuous medial myosin oscillations near the pit.** Time-lapse movie of an exemplary embryo expressing SqhGFP to label myosin II. SqhGFP is shown in the left panel, and the calculated apical-medial myosin II intensity on the right.

**Supplemental Movie 3, related to Figure 3. Extended wild-type embryo movie 1 for strain rate analysis.** Control embryo (*sqh[AX3];;sqhGFP, UbiRFP*) with cell membrane shown in the left panel and myosin II in the right panel. This is ExpID0618.

**Supplemental Movie 4, related to Figure 3. Extended wild-type embryo movie 2 for strain rate analysis.** Control embryo (*sqh[AX3];;sqhGFP, UbiRFP*) with cell membrane shown in the left panel and myosin II in the right panel. This is ExpID0690.

**Supplemental Movie 5, related to Figure 3. Extended *hkb^-/-^* mutant embryo movie 2 for strain rate analysis.** *hkb^-/-^* mutant embryo (*sqhGFP42, UbiRFP; hkb[2]*) with cell membrane shown in the left panel and myosin II in the right panel. This is ExpID0678.

**Supplemental Movie 6, related to Figure 3. Extended *hkb^-/-^* mutant embryo movie 2 for strain rate analysis.** *hkb^-/-^* mutant embryo (*sqhGFP42, UbiRFP; hkb[2]*) with cell membrane shown in the left panel and myosin II in the right panel. This is ExpID0689.

**Supplemental Movie 7, related to Figure 5. Hkb dynamics in the placode.** Embryo of the genotype *UbiRFP, Venus-Hkb* to allow dynamic analysis of Hkb protein levels.

**Supplemental Movie 8, related to Figure 5. Fkh dynamics in the placode.** Embryo of the genotype *UbiRFP, Fkh-GFP* to allow dynamic analysis of Fkh protein levels.

## References

1. Gilmour D, Rembold M, Leptin M. From morphogen to morphogenesis and back. Nature. 2017;541(7637):311–20.

2. Sidor C, Röper K. Genetic Control of Salivary Gland Tubulogenesis in Drosophila. In: Castelli-Gair Hombría J, Bovolenta P, editors. Organogenetic Gene Networks. Switzerland: Srpinger International Publishing; 2016. p. 125–49.

3. Collinet C, Rauzi M, Lenne PF, Lecuit T. Local and tissue-scale forces drive oriented junction growth during tissue extension. Nat Cell Biol. 2015;17(10):1247–58.

4. Lye CM, Blanchard GB, Naylor HW, Muresan L, Huisken J, Adams RJ, et al. Mechanical Coupling between Endoderm Invagination and Axis Extension in Drosophila. PLoS biology. 2015;13(11):e1002292.

5. Chanet S, Miller CJ, Vaishnav ED, Ermentrout B, Davidson LA, Martin AC. Actomyosin meshwork mechanosensing enables tissue shape to orient cell force. Nature communications. 2017;8:15014.

6. Sanchez-Corrales YE, Blanchard GB, Roper K. Radially patterned cell behaviours during tube budding from an epithelium. eLife. 2018;7.

7. Booth AJ, Blanchard GB, Adams RJ, Röper K. A dynamic microtubule cytoskeleton directs medial actomyosin function during tube formation. Dev Cell. 2014;29(5):562–76.

8. Myat MM, Andrew DJ. Organ shape in the Drosophila salivary gland is controlled by regulated, sequential internalization of the primordia. Development. 2000;127(4):679–91.

9. Myat MM, Andrew DJ. Epithelial tube morphology is determined by the polarized growth and delivery of apical membrane. Cell. 2002;111(6):879–91.

10. Myat MM, Andrew DJ. Fork head prevents apoptosis and promotes cell shape change during formation of the Drosophila salivary glands. Development. 2000;127(19):4217–26.

11. Chandrasekaran V, Beckendorf SK. Tec29 controls actin remodeling and endoreplication during invagination of the Drosophila embryonic salivary glands. Development. 2005; 132(15):3515–24.

12. Girdler GC, Röper K. Controlling cell shape changes during salivary gland tube formation in Drosophila. Semin Cell Dev Biol. 2014;31:74–81.

13. Blanchard GB, Kabla AJ, Schultz NL, Butler LC, Sanson B, Gorfinkiel N, et al. Tissue tectonics: morphogenetic strain rates, cell shape change and intercalation. Nat Methods. 2009;6(6):458–64.

14. Spokony R, White K. Spokony insertions, Personal communication to FlyBase. 2013.

15. Chung S, Kim S, Andrew DJ. Uncoupling apical constriction from tissue invagination. eLife. 2017;6:e22235.

16. Dawes-Hoang RE, Parmar KM, Christiansen AE, Phelps CB, Brand AH, Wieschaus EF. folded gastrulation, cell shape change and the control of myosin localization. Development. 2005; 132(18):4165–78.

17. Nikolaidou KK, Barrett K. A Rho GTPase signaling pathway is used reiteratively in epithelial folding and potentially selects the outcome of Rho activation. Curr Biol. 2004;14(20):1822–6.

18. Panzer S, Weigel D, Beckendorf SK. Organogenesis in Drosophila melanogaster: embryonic salivary gland determination is controlled by homeotic and dorsoventral patterning genes. Development. 1992;114(1):49–57.

19. Clark E. Dynamic patterning by the Drosophila pair-rule network reconciles long-germ and short-germ segmentation. PLoS biology. 2017;15(9):e2002439.

20. Röper K. Anisotropy of Crumbs and aPKC Drives Myosin Cable Assembly during Tube Formation. Dev Cell. 2012;23(5):939–53.

21. Kolesnikov T, Beckendorf SK. 18 wheeler regulates apical constriction of salivary gland cells via the Rho-GTPase-signaling pathway. Dev Biol. 2007;307(1):53–61.

22. Morize P, Christiansen AE, Costa M, Parks S, Wieschaus E. Hyperactivation of the folded gastrulation pathway induces specific cell shape changes. Development. 1998;125(4):589–97.

23. Weigel D, Jurgens G, Klingler M, Jackle H. Two gap genes mediate maternal terminal pattern information in Drosophila. Science. 1990;248(4954):495–8.

24. Kerridge S, Munjal A, Philippe JM, Jha A, de las Bayonas AG, Saurin AJ, et al. Modular activation of Rho1 by GPCR signalling imparts polarized myosin II activation during morphogenesis. Nat Cell Biol. 2016;18(3):261–70.

25. Manning AJ, Peters KA, Peifer M, Rogers SL. Regulation of epithelial morphogenesis by the G protein-coupled receptor mist and its ligand fog. Sci Signal. 2013;6(301):ra98.

26. Hanlon CD, Andrew DJ. Outside-in signaling--a brief review of GPCR signaling with a focus on the Drosophila GPCR family. J Cell Sci. 2015;128(19):3533–42.

27. Maybeck V, Röper K. A targeted gain-of-function screen identifies genes affecting salivary gland morphogenesis/tubulogenesis in Drosophila. Genetics. 2009;181(2):543–65.

28. Lim B, Levine M, Yamazaki Y. Transcriptional Pre-patterning of Drosophila Gastrulation. Curr Biol. 2017;27(4):610.

29. Heer NC, Miller PW, Chanet S, Stoop N, Dunkel J, Martin AC. Actomyosin-based tissue folding requires a multicellular myosin gradient. Development. 2017;144(10):1876–86.

30. Bailles A, Collinet C, Philippe JM, Lenne PF, Munro E, Lecuit T. Genetic induction and mechanochemical propagation of a morphogenetic wave. Nature. 2019;572(7770):467–73.

31. Sidor C, Stevens TJ, Jin L, Boulanger J, Roper K. Rho-Kinase Planar Polarization at Tissue Boundaries Depends on Phospho-regulation of Membrane Residence Time. Dev Cell. 2020;52(3):364–78 e7.

32. Simoes S, Denholm B, Azevedo D, Sotillos S, Martin P, Skaer H, et al. Compartmentalisation of Rho regulators directs cell invagination during tissue morphogenesis. Development. 2006;133(21):4257–67.

33. Lye CM, Naylor HW, Sanson B. Subcellular localisations of the CPTI collection of YFP- tagged proteins in Drosophila embryos. Development. 2014;141(20):4006–17.

34. Royou A, Field C, Sisson JC, Sullivan W, Karess R. Reassessing the role and dynamics of nonmuscle myosin II during furrow formation in early Drosophila embryos. Mol Biol Cell. 2004;15(2):838–50.

35. Henderson KD, Andrew DJ. Regulation and function of Scr, exd, and hth in the Drosophila salivary gland. Dev Biol. 2000;217(2):362–74.

36. Zhou B, Bagri A, Beckendorf SK. Salivary gland determination in Drosophila: a salivary-specific, fork head enhancer integrates spatial pattern and allows fork head autoregulation. Dev Biol. 2001;237(1):54–67.

37. Förster D, Luschnig S. Src42A-dependent polarized cell shape changes mediate epithelial tube elongation in Drosophila. Nat Cell Biol. 2012;14(5):526–34.

38. Hampel S, Chung P, McKellar CE, Hall D, Looger LL, Simpson JH. Drosophila Brainbow: a recombinase-based fluorescence labeling technique to subdivide neural expression patterns. Nat Methods. 2011;8(3):253–9.

39. McDonald JA, Doe CQ. Establishing neuroblast-specific gene expression in the Drosophila CNS: huckebein is activated by Wingless and Hedgehog and repressed by Engrailed and Gooseberry. Development. 1997;124(5):1079–87.

40. Weigel D, Bellen HJ, Jurgens G, Jackle H. Primordium specific requirement of the homeotic gene fork head in the developing gut of the Drosophila embryo. Roux Arch Dev Biol. 1989;198(4):201–10.

41. Fuse N, Yu F, Hirose S. Gprk2 adjusts Fog signaling to organize cell movements in Drosophila gastrulation. Development. 2013;140(20):4246–55.

42. Boulanger J, Kervrann C, Bouthemy P, Elbau P, Sibarita JB, Salamero J. Patch-based nonlocal functional for denoising fluorescence microscopy image sequences. IEEE Trans Med Imaging. 2010;29(2):442–54.

43. Blanchard GB. Taking the strain: quantifying the contributions of all cell behaviours to changes in epithelial shape. Philosophical transactions of the Royal Society of London Series B, Biological sciences. 2017;372(1720).

44. Butler LC, Blanchard GB, Kabla AJ, Lawrence NJ, Welchman DP, Mahadevan L, et al. Cell shape changes indicate a role for extrinsic tensile forces in Drosophila germ-band extension. Nat Cell Biol. 2009;11(7):859–64.

45. Blanchard GB, Murugesu S, Adams RJ, Martinez-Arias A, Gorfinkiel N. Cytoskeletal dynamics and supracellular organisation of cell shape fluctuations during dorsal closure. Development. 2010;137(16):2743–52.

